# Fecal microbial transfer and complex carbohydrates mediate protection against COPD

**DOI:** 10.1101/2023.10.16.562613

**Authors:** Kurtis F. Budden, Shakti D. Shukla, Kate L. Bowerman, Shaan Gellatly, David L.A. Wood, Nancy Lachner, Sobia Idrees, Vyoma K. Patel, Alen Faiz, Saima Firdous Rehman, Chantal Donovan, Charlotte A. Alemao, SJ Shen, Kanth S. Vanka, Jazz Mason, Tatt Jhong Haw, Michael Fricker, Simon Keely, Nicole G. Hansbro, Gabrielle T. Belz, Jay C. Horvat, Thomas M. Ashhurst, Caryn van Vreden, Helen M. McGuire, Barbara Fazekas de St Groth, Nicholas J.C. King, Ben Crossett, Stuart J. Cordwel, Lorenzo Bonaguro, Joachim L. Schultze, Samuel C Forster, Matthew A. Cooper, Leopoldo N. Segal, Annalicia Vaughan, Peter F. Collins, Rayleen V. Bowman, Kwun M. Fong, Ian A. Yang, Peter A. Wark, Paul G. Dennis, Philip Hugenholtz, Philip M. Hansbro

## Abstract

**Objective:** Chronic obstructive pulmonary disease (COPD) is a major cause of global illness and death, most commonly caused by cigarette smoke. The mechanisms of pathogenesis remain poorly understood, limiting the development of effective therapies. The gastrointestinal microbiome has been implicated in chronic lung diseases *via* the gut-lung axis, but its role is unclear.

**Design:** Using an *in vivo* mouse model of cigarette smoke-induced COPD and fecal microbial transfer (FMT), we characterized the fecal microbiota using metagenomics, proteomics and metabolomics. Findings were correlated with airway and systemic inflammation, lung and gut histopathology, and lung function. Complex carbohydrates were assessed in mice using a high resistant starch diet, and in sixteen COPD patients using a randomized, double-blind, placebo-controlled pilot study of inulin supplementation.

**Results:** FMT alleviated hallmark features of COPD (inflammation, alveolar destruction, impaired lung function), gastrointestinal pathology and systemic immune changes. Protective effects were additive to smoking cessation. Disease features correlated with the relative abundance of *Muribaculaceae, Desulfovibrionaceae* and *Lachnospiraceae* family members. Proteomics and metabolomics identified downregulation of glucose and starch metabolism in cigarette smoke-associated microbiota, and supplementation of mice or human patients with complex carbohydrates improved disease outcomes.

**Conclusion:** The gut microbiome contributes to COPD pathogenesis and can be targeted therapeutically.

**What is already known on this topic:**

- Changes in gut microbiota are associated with COPD but the underlying host and microbial mechanisms are unclear, limiting the therapeutic applications.

**What this study adds:**

- Microbiome composition and metabolism is reproducibly correlated with lung and gastrointestinal pathology in experimental COPD.
- Microbiome modifying interventions effectively alleviate disease, including protective effects supplementing smoking cessation.
- Nutritional interventions targeting the microbiome in COPD patients demonstrate efficacy in a small pilot study.

**How this study might affect research, practice or policy:**

- Microbiome-targeting therapeutics and nutritional interventions may be developed for COPD, including as supplements to smoking cessation.

## MAIN TEXT

Chronic obstructive pulmonary disease (COPD) is most often caused by long-term cigarette smoke (CS) inhalation and includes chronic inflammation, airway remodeling, and emphysema leading to progressive lung function impairment which persists after smoking cessation.^1^ Over 210 million people globally have COPD, causing >3.3 million deaths annually although even these are substantially underreported.^2^ Pharmacological treatments have little efficacy in reversing disease, suppressing its progression, or preventing mortality and have significant adverse effects.^3^ Mechanisms of COPD pathogenesis remain poorly understood and no singular driver of disease has been discovered, suggesting that chronic effects from a range of factors drive disease pathogenesis over many years.^4^ Development of effective therapies will require targeting this array of factors.

The microbiome is under intense investigation for its immunoregulatory capacity and association with disease.^5^ Most COPD studies have focused on respiratory microbiota, demonstrating increased bioburden, reduced diversity, and enrichment of *Firmicutes* and *Proteobacteria*.^5^

The gut hosts the largest and most diverse microbiome of the human body that, depending upon its composition, can drive or suppress inflammation, including in the lung.^6^ Alterations in the gut microbiome from antibiotics or diet influence asthma development and respiratory infections.^6^ Chronic CS-exposure induces gastrointestinal histopathology,^7^ and COPD patients have increased risk of inflammatory bowel diseases^6^ and altered gut microbiome composition.^8^

Transfer of whole microbial communities from healthy individuals through fecal microbial transfer (FMT) is an effective therapy in patients with *Clostridioides difficile* colitis, but its implementation in other diseases is not as well supported.^9^ FMT may therefore have cost-effective benefits in COPD, but further examination is required. A study using a short-term smoke and poly I:C (TLR3 agonist) model showed that FMT prevented emphysema and identified associations with *Bacteroidaceae* and *Lachnospiraceae* families by 16S gene sequencing,^10^ but important controls and analyses linking taxa to pathogenesis were lacking and the model was not representative of human CS-induced COPD. Similarly, *Parabacteroides goldsteinii* lipopolysaccharide (LPS) had protective effects in a murine model of COPD, but broader associations between microbiota and disease were not identified.^11^ Crucially, neither study identified bacteria associated with COPD in human studies^8^ or demonstrated human translation.

We hypothesized that gastrointestinal microbiota contribute to COPD pathogenesis, and provide a detailed characterization of the gastrointestinal microbiome in experimental CS-induced COPD using multi-omics. FMT protected against experimental COPD through changes in microbiota that correlated with key disease features. Proteomics and metabolomics implicated restoration of bacterial complex carbohydrate metabolism in the protective effects, supported by interventional studies in experimental COPD and human COPD patients.

## Methods

### Mice, CS-exposure, microbiome transfer and diet studies

Female C57BL/6 mice (3-5 weeks old) from the University of Newcastle Animal service Unit (Newcastle, Australia) underwent three weeks of baseline microbiome normalization by transferring soiled bedding and co-housing. Microbiome normalization was not performed for validation experiments (8 weeks CS).

Mice were exposed to normal room air or CS from twelve 3R4F cigarettes (University of Kentucky, Lexington, KY) twice per day, 5 days per week, for 8 or 12 weeks as previously described.^12–20^ FMT was administered by transfer of soiled bedding or oral gavage of faecal supernatants twice per week. For diet studies, mice were fed a conventional semi-pure diet (AIN93G) or a resistant starch diet (SF11-025; Specialty Feeds, WA, Australia) *ad libitum* commencing 2 week prior to CS-exposure and maintained until the end of experiment. Feces were collected weekly, with at least 3 fresh fecal pellets collected and stored at −80°C until processing. Airway inflammation, lung and colon histopathology, lung function and gene expression analyses were assessed as previously described.^7, 12–21^ Detailed methods are provided in the online supplement.

All experiments were approved by University of Newcastle Animal Ethics Committee.

### Analysis of gut microbiome

Faecal microbiome composition, including metagenomics and 16S rRNA gene sequencing, were analysed as previously described^8^ with detailed methods provided in the online supplement. Briefly, α-diversity was calculated using QIIME v1.8.0^22^, principal component analysis performed using the R package vegan v2.5-1^23^ within metagenomeSeq v1.22.0,^24^ and differential abundance determined using DESeq2 with Benjamini-Hochberg adjustment. Sparse Partial Least-Squares Discriminant Analysis (sPLS-DA) was conducted using the R package mixOmics v6.3.2^25^. Host phenotypes were tested for association with microbiome composition using the envfit function within the vegan R package. Spearman’s rho was calculated using the ‘corr.test’ function within the R package psych v1.8.12.^26^

### CyTOF analysis

For time-of-flight mass cytometry, single cell suspensions of bone marrow, blood, and spleen cells were stained with cell cycle marker iododeoxyuridine (IdU), viability marker cisplatin, and surface and intracellular antibodies (Table S17). Expression levels of 38 markers were measured using a Helios instrument (Fluidigm), transformed with a logicle transformation^27^ and used for Uniform Manifold Approximation and Projection for Dimension Reduction (UMAP) using the naïve R implementation (umap 0.2.7). Cells clusters were calculated using Rphenograph (Github JinmiaoChenLab v. 0.99.1)^28^. Detailed methods are provided in the online supplement.

### Cell culture

Raw 264.7 monocytes (1×10^6^) were seeded into 12-well plates. Cells were incubated in media alone, or 1:1,000 dilutions of sterile-filtered fecal homogenate (100mg/mL) from CS- or air-exposed mice for 24 hours. During the final 4 hours, lipopolysaccharide (1µg/mL), monophosphoryl lipid A (4µg/mL), lipoteichoic acid (1µg/mL) were added to cell media. TNFα protein in culture media was assessed using Duoset ELISA kits (R & D Systems).

### Proteomics of mouse feces

Fecal proteins were analysed by nanoflow liquid chromatography (Ultimate 3000RSLCnano, Thermo Scientific) with peptides introduced via an EasySpray Nano source coupled to a Q-Exactive Plus Quadrupole Orbitrap mass spectrometer (Thermo Scientific) and subjected to data dependent tandem mass spectrometry (MS/MS). .raw files were processed in Proteome Discoverer v2.1 using the Sequest HT algorithm^29^, searched against the mouse gut microbiota GigaDB database,^30^ and integrated with metagenomics results using the psych package in R^31^ and Diablo mixOmics package. Detailed methods are provided in the online supplement.

### Metabolomics

Metabolomics of caecum contents was performed by Metabolon Inc (Durham, NC) using the Global HD4 mass spectrometry platform as previously described^8^. Detailed methods are provided in the online supplement.

### Human interventional study

Sixteen COPD patients were recruited from June 2019 to March 2020 at the Respiratory Investigation Unit at The Prince Charles Hospital (TPCH) with written and informed consent (approved by the TPCH Human Research Ethics Committee – HREC/18/QPCH/234). Nine participants were randomized into the intervention arm (inulin 10g daily for 4 weeks) and seven to the placebo group (maltodextrin 10g daily for 4 weeks). Age, gender, body mass index (BMI), smoking history (pack years) and spirometry (forced expiratory volume in 1 second/forced vital capacity [FEV_1_/FVC] and FEV1% predicted) were similar between the two groups (Table 1).

**Table 1:** Patient characteristics.

|  | Placebo (n=7) |  | Inulin (n=9) |  | P-value |
| --- | --- | --- | --- | --- | --- |
|  | Mean | SD | Mean | SD |  |
| Age | 69.71 | 6.50 | 72.78 | 6.46 | 0.18 |
| Gender (M, %) | 42.86 | N/A | 66.67 | N/A | 0.34 |
| BMI | 29.68 | 6.06 | 28.66 | 4.85 | 0.71 |
| Smoking pack years | 56.21 | 20.02 | 49.73 | 24.94 | 0.6 |
| FEV1/FVC | 0.48 | 0.09 | 0.48 | 0.13 | 0.95 |
| FEV1% | 55.43 | 18.04 | 54.56 | 16.18 | 0.98 |
| Freq. exacerbators (n, %) | 5 (71%) | N/A | 3 (33%) | N/A | 0.13 |
| **Medications (n, %)** |  |  |  |  |  |
| Inhaled corticosteroids (ICS) | 5 (71%) | N/A | 7 (77%) | N/A | 0.77 |
| Beta agonists | 7 (100%) | N/A | 9 (100%) | N/A | >0.99 |
| LAMA | 1 (14%) | N/A | 2 (22%) | N/A | 0.69 |
| Statin | 2 (28%) | N/A | 6 (67%) | N/A | 0.13 |
| PPI | 4 (57%) | N/A | 3 (33%) | N/A | 0.34 |
| ACE | 2 (28%) | N/A | 2 (22%) | N/A | 0.77 |
| Beta blockers | 1 (14%) | N/A | 1 (11%) | N/A | 0.85 |
| Angiotensin II receptor antagonist | 2 (28%) | N/A | 1 (11%) | N/A | 0.37 |
| SSRI/SNRI | 2 (28%) | N/A | 1 (11%) | N/A | 0.37 |
| **Season during intervention (n, %)** |  |  |  |  |  |
| Spring/summer | 2 (28%) | N/A | 3 (33%) | N/A | 0.84 |

### Statistical Analysis

Except where specified, data was analyzed in GraphPad Prism v9.0 (San Diego, California). Outliers were identified by Grubbs test, and excluded from analysis only if investigators had reported technical errors during sample collection/processing before analysis. Data was analyzed by one-way ANOVA with Holm-Šídák’s post hoc test or Student’s T test, or non-parametric equivalents, for data from mice. Data from the COPD patient cohort was analyzed by Mann-Whitney test (continuous variables) or Chi^2^ test (categorical variables).

## Results

### FMT alleviated dysbiosis and disease features in experimental COPD

To define changes in microbiome composition and the effects of FMT, C57BL/6 mice underwent microbiome normalization to control for variability in starting microbiome composition before exposure to mainstream CS through the nose only for twelve weeks (12 wk CS),^12–15, 18–20^ with a subset of mice modeling smoking cessation with eight weeks of CS-exposure followed by four weeks of rest (8 wk CS + 4 wk rest). Mice were treated *via* passive FMT through transfer of soiled bedding from air-exposed mice, with controls maintained in their own bedding.

Shotgun metagenomics of fecal samples collected prior to interventions (week 0) and at completion (week 12) recovered seventy-four metagenome-assembled genomes (MAG) >80% complete (dereplicated at 95% identity) representing 12 families. Microbiome composition was assessed with public genomes within Genome Taxonomy Database (GTDB) release 03-RS83.^32^ All experimental groups experienced a shift in microbiome composition associated with the maturation of the mice between week 0 and 12 (from 6 to 18 weeks old), with increased α-diversity enrichment of species belonging to the genus *Prevotella* and family *Muribaculaceae*, and depletion of *Akkermansia* and *Duncaniella* (a member of *Muribaculaceae*) species (Fig. S1A-C, Table S5).

While different experimental groups had similar α-diversity at week 12, microbiome composition differed significantly (Fig. S1d-e; P=0.01, PERMANOVA of Bray-Curtis distances). CS-exposure was associated with increased *Phocaeicola vulgatus*, *Muribaculaceae* species *Duncaniella sp001689575* and *Amulumruptor sp001689515*, *Desulfovibrionaceae* species *Mailhella sp003512875* and *Akkermansia muciniphila*, and decreased *Lachnospiraceae* species *Muricomes sp001517425* (Fig. 1a-b, Table S6-7). CS-exposed FMT mice maintained increased abundance of most CS-associated species compared to air-exposed mice, except for the *Lachnospiraceae* member *UBA3282 sp009774575* (MAG FTS36, Table S6). This species, along with CS-associated *Mailhella sp003512875* (MAG FTS70), was identified in a multi-variate analysis-derived signature distinguishing 12-week CS-exposed mice from both air-and 12-week CS-exposed FMT mice (Fig. 1c-d, Table S8). Both species were increased with CS-exposure and decreased with FMT, consistent with a role in disease (Fig. 1e).

**Figure 1.**
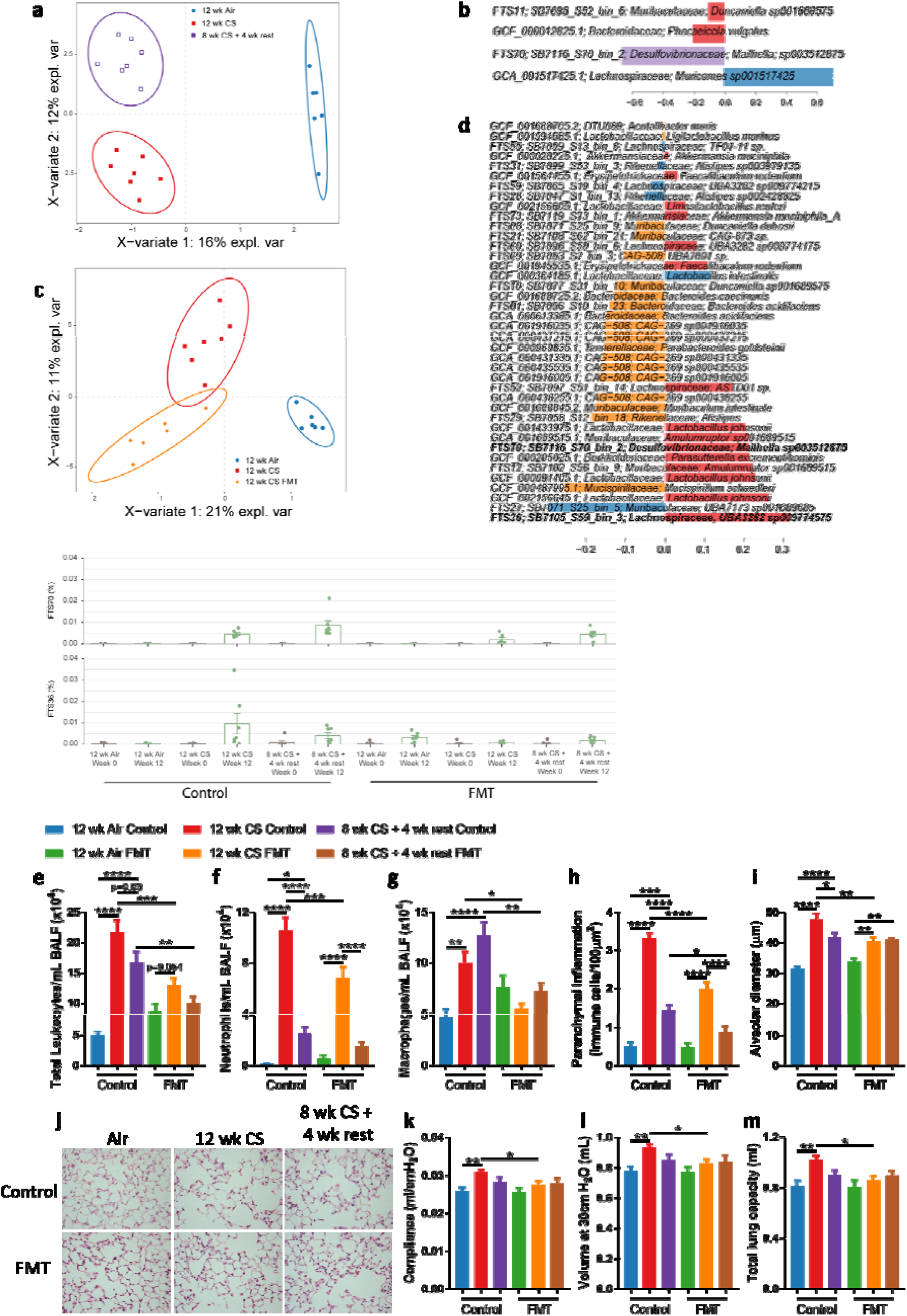
Fecal microbiota transfer (FMT) alleviated dysbiosis and disease features in severe experimental COPD and smoking cessation. Mice were exposed to CS or normal air for 12 weeks, or 8 weeks CS followed by 4 weeks with normal air (8 wk CS + 4 wk rest). Mice also received FMT through transfer of soiled bedding or were maintained in their own bedding (Control) for 12 weeks. **a-d,** Fecal samples were collected at the end of experiment (week 12) and analyzed using shotgun metagenomics. (**a**) Multi-variate analysis (sPLS-DA) of CS- and air-exposed mice at week 12 demonstrating distinction between the groups along component 1 based on relative abundance at the genome level, centered log-ratio transformed and (**b**) species contributing to that distinction. (**c**) Multi-variate analysis (sPLS-DA) of 12-week CS-exposed control and FMT treated mice, and air-exposed mice, demonstrating distinction between CS-exposed control and FMT mice along component 2 and (**d**) species contributing to that distinction. (**e**) Relative abundance of *Lachnospiraceae* FTS36 and *Mailhella sp003512875* FTS70 were increased with CS-exposure, which were alleviated by FMT. **f-n,** hallmark features of COPD assessed at the completion of the experiment (week 12). FMT alleviated CS-induced increases in (**f-h**) total leukocytes, neutrophils and macrophages in bronchoalveolar lavage fluid (BALF), and (**i**) parenchymal inflammation in 12-week CS and 8-week CS + 4-week rest mice. FMT also alleviated CS-induced increases in (**j-k**) emphysema-like alveolar enlargement, and (**l-n**) lung function parameters of compliance, volume and total lung capacity in 12-week CS mice. N = 8 per group (**a-n**). Data presented as mean +/- standard error of the mean. * = p<0.05; ** = p<0.01; *** = p<0.001; **** = p<0.0001 using one-way ANOVA with Holm-Sidak’s post-hoc analysis (**f-n**).

CS-exposure induced lung inflammation characterized by increased total leukocytes, neutrophils and macrophages in BALF, and immune cells in parenchyma (Fig. 1g-i), as well as emphysematous alveolar destruction (Fig. 1j-k). BALF macrophages remained elevated after smoking cessation at levels comparable to mice exposed to CS for 12 weeks, while other measures of inflammation and alveolar destruction were present but had reduced severity. Impaired lung function, with increased compliance, volume and total lung capacity, was observed after 12 weeks CS-exposure (Fig. 1l-n).

FMT-treated mice had significantly lower BALF and parenchymal inflammation in mice exposed to CS for 12 weeks, or with smoking cessation (Fig. 1e-h). Unlike smoking cessation, FMT reduced BALF macrophages, and the combination of smoking cessation and FMT had additive effects by further reducing total leukocytes and parenchymal inflammation. Most importantly, FMT alleviated both emphysema and impaired lung function after CS for 12 weeks (Fig. 1i-m).

Thus, FMT alleviated hallmark features of COPD and improved the resolution of chronic inflammation with smoking cessation.

### FMT alters systemic manifestations of COPD

Given our evidence of CS-induced systemic co-morbidities,^7, 12^ we assessed gut manifestations of COPD and systemic leukocyte populations in mice exposed to CS for 12 weeks, with and without FMT. CS-induced increases in colonic sub-mucosal fragmentation and vascularization, and colon tissue mRNA expression of the microbe-sensing pattern recognition receptors *Tlr3* and *Tlr4* was prevented by FMT, with similar trends for *Tlr2* (Fig. 2a-e). No significant differences were observed in expression of *Tlr9* (Fig. 2f).

**Figure 2.**
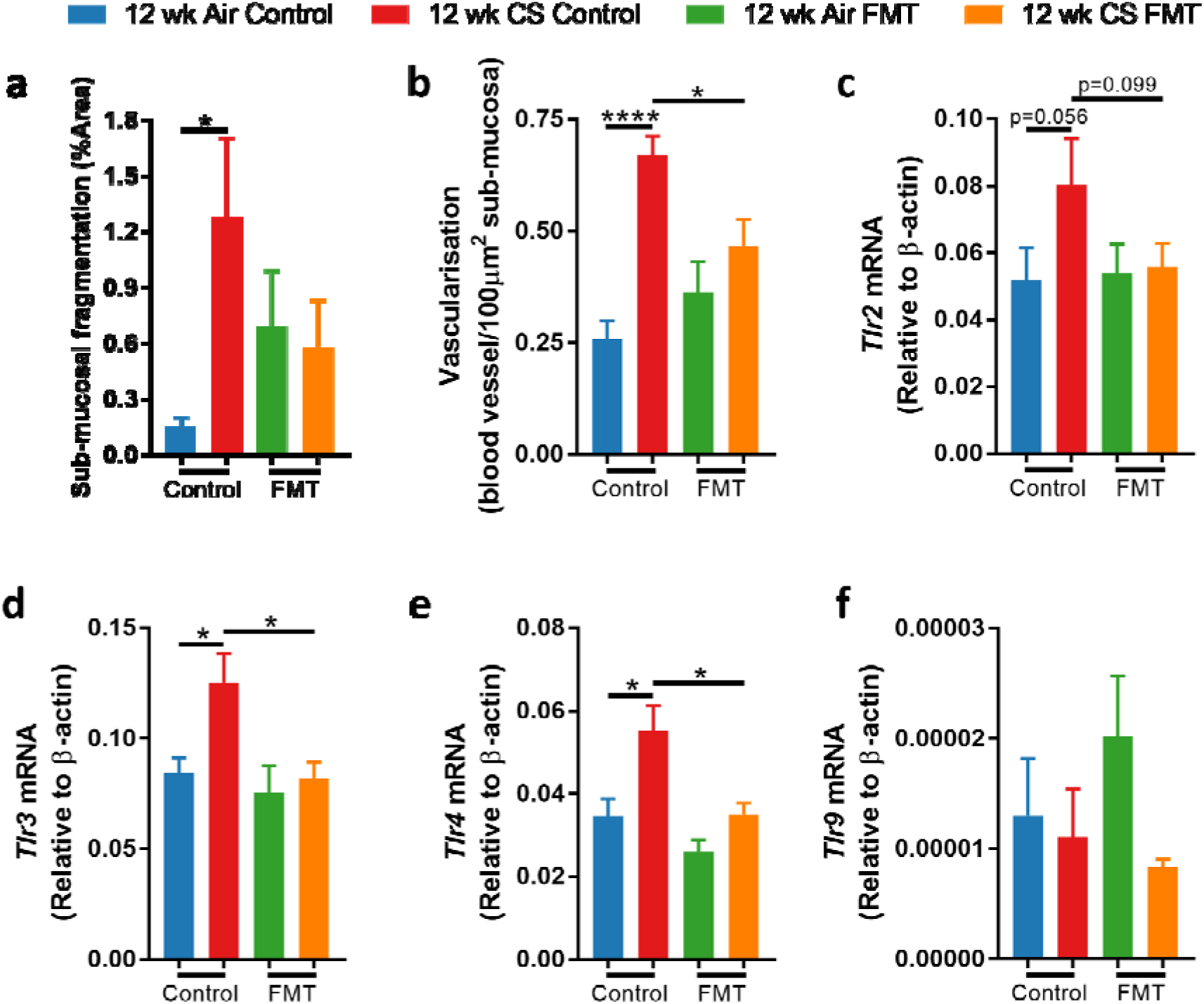
Fecal microbial transfer (FMT) alleviated cigarette smoke (CS)-induced colon histopathology and microbial sensor expression changes. **a-f,** Mice were exposed to CS or normal air for 12 weeks, and received FMT through transfer of soiled bedding or were maintained in their own bedding (Control). FMT alleviated CS-induced increases in (**a**) sub-mucosal fragmentation, (**b**) vascularization and expression of (**c**) *Tlr2* (non-significant), (**d**) *Tlr3*, and (**e**) *Tlr4* in colon tissue. (**f**) Expression of *Tlr9* was not altered by either CS or FMT. N = 7-8 per group. Data presented as mean +/- standard error of the mean. * = p<0.05 using one-way ANOVA with Holm-Sidak’s post-hoc analysis.

We employed deep immune profiling using cytometry by time-of-flight (CyTOF) with 38 immune cell markers in the bone marrow, blood and spleen. Dimensionality reduction and clustering of bone marrow cells identified 20 clusters with coherent protein expression, but few were altered (Fig. 3a-c). CS-exposure reduced the abundance of B cells, likely driven by a downregulation of proliferating (IdU^+^) B cells, but these effects were not alleviated by FMT (Fig. 3d). In blood, 25 clusters were identified (Fig. 3e-g). B cells were reduced by CS-exposure and partially restored by FMT (Fig 3h). CS-exposure also increased blood Ly6C^lo^ monocytes, and FMT restored their abundance to levels in air-exposed mice (Fig. 3h). In the spleen, 23 clusters were identified (Fig. 3i-k) with CD8^+^ conventional dendritic cells (cDCs) and B cells reduced while CD107^+^ progenitors and Ly6C^lo^ monocytes were increased by CS-exposure (Fig. 3l). FMT increased the abundance of CD8^+^ cDCs, CD107^+^ progenitors and Ly6C^lo^ monocytes, partially restoring the CS-induced depletion of CD8a^+^ cDCs but enhancing the impact of CS-induced increases in CD107^+^ progenitors and Ly6C^lo^ monocytes.

**Figure 3.**
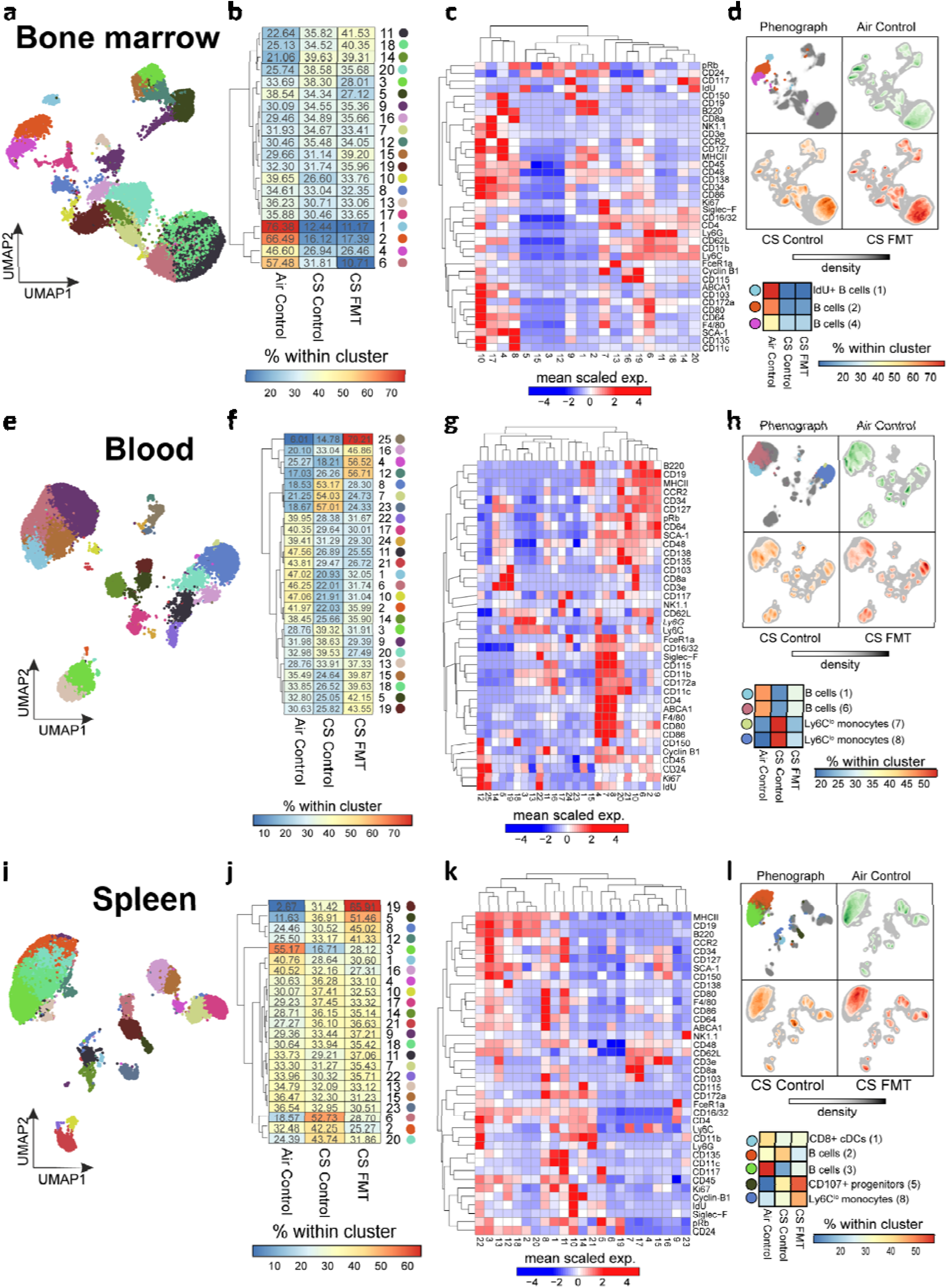
Cigarette smoke (CS)-exposure altered systemic leukocyte populations, and fecal microbial transfer (FMT) alleviated depletion of some blood and splenic populations. **a-l,** Mice were exposed to CS or normal air for 12 weeks and received FMT through transfer of soiled bedding or were maintained in their own bedding (Control). Cell abundances were quantified using cytometry by time-of-flight (CYToF). (**a, b**) Dimensionality reduction and clustering of bone marrow cells identified 20 clusters of cells (**c**) based upon the scaled mean marker expression. (**d**) Uniform manifold approximation and projection (UMAP) and a confusion matrix of the normalized frequencies of clusters demonstrated CS-induced depletion of B cells and IDU^+^ B cells which was not alleviated by FMT. (**e, f**) Dimensionality reduction and clustering of blood cells identified 20 clusters of cells (**g**) based upon the scaled mean marker expression. (**h**) UMAP and a confusion matrix of the normalized frequencies of clusters demonstrated a CS-induced depletion of B cells and increase of Ly6C^lo^ monocytes, which were alleviated by FMT. (**i, j**) Dimensionality reduction and clustering of splenic cells identified 23 clusters of cells (**k**) based upon the scaled mean marker expression. (**l**) UMAP and a confusion matrix of the normalized frequencies of clusters demonstrated CS-induced depletion of CD8^+^ conventional dendritic cells (cDCs) and B cells, and a CS-induced increase of CD107^+^ progenitors and Ly6C^lo^ monocytes. FMT increased the abundance of CD8^+^ cDCs, CD107^+^ progenitors and Ly6C^lo^ monocytes compared to CS control mice, partially reducing CD8^+^ cDCs but enhancing the CS-induced increases in CD107^+^ progenitors and Ly6C^lo^ monocytes. N = 6 per group.

Thus, FMT alleviated CS-induced colonic pathology and TLR expression, as well as CS-induced depletion of blood B cells and splenic CD8^+^ cDCs and increases in blood Ly6C^lo^ monocytes.

### Correlations of lung and gut pathology with microbiota

We assessed the role of the gut microbiome in pathogenesis by fitting phenotypic data to a β-diversity ordination analysis, identifying significant associations with lymphocytes and emphysema consistent with CS-induced separation in microbiome composition (Fig. 4a). When smoking cessation groups were excluded to assess associations in more severe disease, total leukocytes and neutrophils were also associated with microbiome composition (Fig. 4b). Correlations between individual species and phenotypes identified positive correlations between *Mailhella sp003512875* (FTS70) and inflammation, emphysema, colon vascularization and *Tlr4* expression (Fig. 4c, Table S9). *Muribaculaceae* species *Amulumruptor sp001689515* and *A. muciniphila* positively correlated with lymphocytes and emphysema while *A. muciniphila_A* correlated with lymphocytes and colon *Tlr3* expression. *Muribaculum intestinale* and *Muribaculaceae* species *UBA7173 sp001689685* were negatively correlated with inflammation, emphysema and colon *Tlr4* expression indicating a protective role.

**Figure 4.**
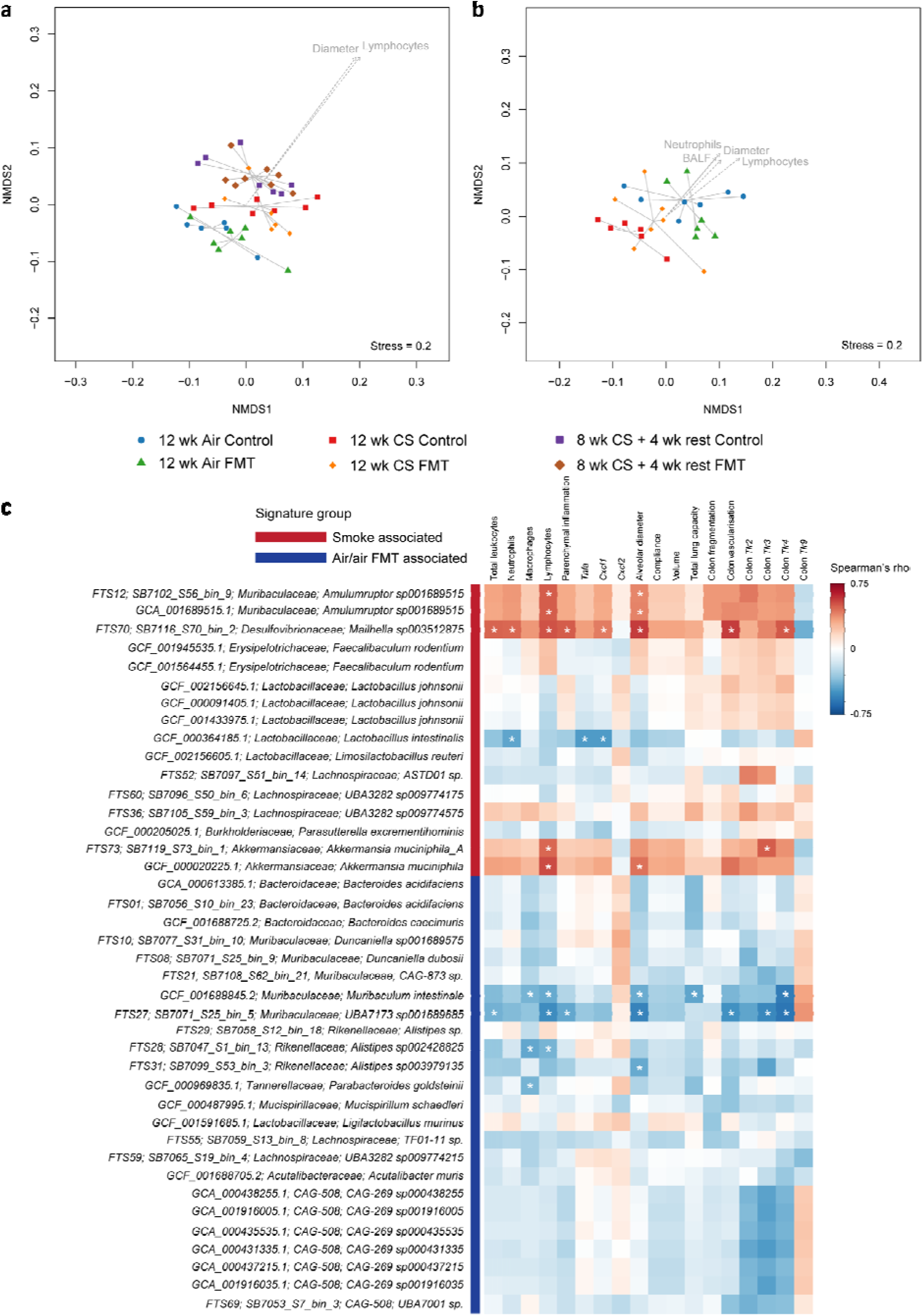
Cigarette smoke (CS)-associated bacterial species positively correlate with phenotypic measurements. **a-c,** Mice were exposed to CS (12 wk CS) or normal air (Air) for 12 weeks, or 8 weeks CS followed by 4 weeks with normal air (8 wk CS + 4 wk rest). Mice also received fecal microbiota transfer (FMT) of soiled bedding or were maintained in their own bedding (Control). (**a, b**) Nonmetric multi-dimensional scaling (NMDS) ordination plot of Bray-Curtis distances with phenotypic variables significantly associated with CS and/or FMT fitted to axes. Lung compliance was excluded due to co-linearity with volume. (**a**) Lymphocytes and alveolar diameter were significantly associated with microbiome composition (Benjamini-Hochberg, P<0.05). (**b**) A similar analysis was performed excluding smoking cessation mice (8 wk + 4 wk rest), which also identified total leukocytes in bronchoalveolar lavage (BALF) and neutrophils. (**c**) Spearman correlation between phenotypic data and relative abundance of genomes separating CS-exposed mice (red bar) from air-exposed mice or CS-exposed mice receiving FMT (blue bar). Squares marked with an internal asterisk have p<0.05, demonstrating key associations with bacteria. N = 8 per group.

### CS-induced changes in microbiota impair responsiveness to TLR4 agonists

Given that CS-associated taxa correlated with colonic TLR expression, we assessed the effects of CS-induced dysbiosis on immune responses *in vitro*. RAW264.7 mouse monocytes were cultured for 24 hours in media alone, or in sterile-filtered homogenates of feces from mice exposed to CS or air for 12 weeks. During the final 4 hours of incubation, TLR4 agonists LPS and monophosphoryl lipid A (MPLA), or the TLR2 agonist lipoteichoic acid (LTA) were added to directly stimulate TLRs. Feces from either CS-or air-exposed mice induced TNFα production compared to media alone, but this was significantly lower with feces from CS-exposed mice (Fig. S2). LPS and MPLA induced TNFα production at comparable levels in cells with media or feces from air-exposed mice, but feces from CS-exposed mice suppressed LPS- and MPLA-induced TNFα. There were no significant differences in LTA-induced TNFα production, suggesting that the impairment may be specific to TLR4.

### Experimental COPD and FMT were consistently associated with bacterial taxa

To validate our findings and explore temporal changes in the microbiome up until disease onset, mice were exposed to CS for eight weeks with passive FMT through transfer of soiled bedding. Changes in gut microbiota were profiled weekly using 16S rRNA gene sequencing of fresh fecal samples collected directly from each mouse for the duration of the experiment. Microbiota composition in different groups were distinguishable from week one and continued to transition over time (Fig. S3a-c). Multi-variate analysis using sPLS-DA revealed CS-associated separation at week eight (Fig. S3d) driven primarily by five amplicon sequence variants (ASVs), including two *Muribaculaceae* family members which increased in CS-exposed mice (without FMT) from week three but only increased in CS-exposed mice receiving FMT after six weeks and decreased in subsequent weeks (Fig. S3e-f).

We also explored microbiome changes in gastrointestinal tissues collected at week 8 and observed similar CS-associated separation in colons, driven by the previously identified *Muribaculaceae* ASVs plus *Desulfovibrionaceae* and *Lachnospiraceae* species (Fig. S4a, d). CS-associated separation was less clear in the cecum (Fig. S4b, e) and was not evident in the ileum (Fig. S4c, f). However, ileum samples did reveal separation between CS-exposed FMT and CS-exposed control mice, with the abundance of distinguishing ASVs in the FMT group resembling air-exposed levels (Fig. S4g-k).

Phenotypic measures were collected at the end of the experiment (Week 8), with CS-induced induced inflammation, alveolar destruction and lung function impairment (Fig. S3g-p). FMT blunted the CS-induced increase in BALF cells and completely suppressed changes in total lung capacity (Fig. S3g-i, p). Similar phenotypic changes were observed with active FMT administered by oral gavage, confirming the involvement of gut microbiota (Fig. S5a-f).

Relative abundance of CS-associated *Muribaculaceae 91ca* in feces at the final timepoint (Week 8) positively correlated with total leukocytes, emphysema, and work of breathing (Fig. S6a; Table S12). ASVs associated with non-smoking had a variable abundance pattern across the experiment and did not correlate with any phenotypic scores (Fig. S6b-d).

Thus CS-induced experimental COPD is reproducibly associated with changes in gut microbiota, particularly a *Muribaculaceae* family member, and FMT alleviated hallmark features of disease.

### Multi-omics implicate downregulation of microbial glucose and starch metabolism in experimental COPD

To assess changes in microbial function, we characterized the fecal proteome (Table S13) and integrated the results with bacterial abundances. Correlations with individual MAGs did not reveal clear associations, but genera *CAG-307*, *Staphylococcus*, *Bilophila*, *Aerococcus*, *CAG-411*, *Corynebacterium*, *Rikenella*, *Lactonifactor*, *Acetivibrio* and *Absiella* were most frequently correlated with the microbial proteome (R≥0.7; p<0.05; Fig. 5a, Table S14).

**Figure 5:**
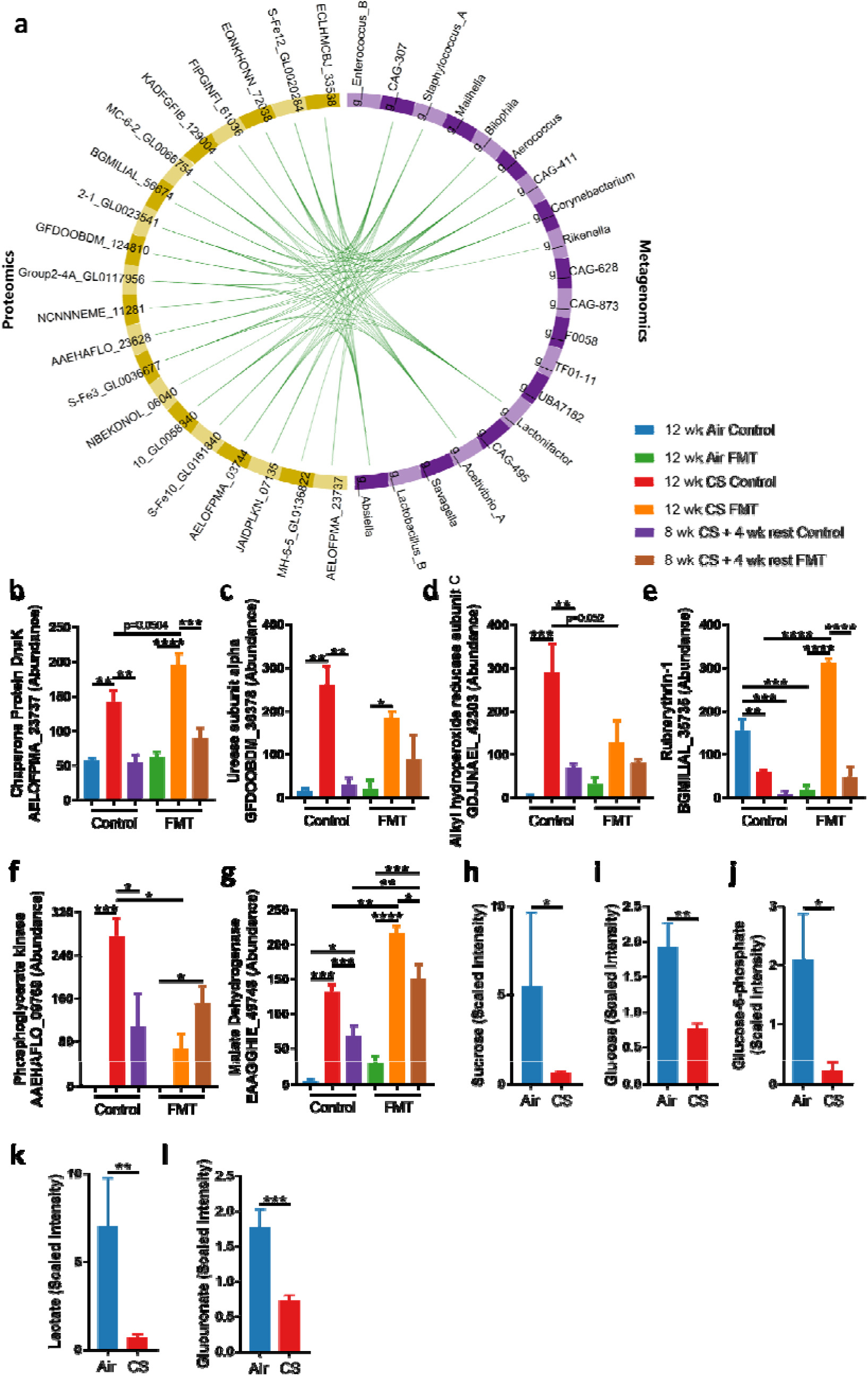
Proteomics and metabolomics show cigarette smoke (CS)- and FMT-induced alterations in oxidative stress responses and glucose metabolism. **A-h,** Mice were exposed to CS (12 wk CS) or normal air (Air) for 12 weeks, or 8 weeks CS followed by 4 weeks with normal air (8 wk + 4 wk rest). Mice also received FMT through transfer of soiled bedding or were maintained in their own bedding (Control) for 12 weeks. (a) Pearson correlation of highly abundant bacterial genera (purple) from metagenomics and proteins (yellow) from proteomics using the MixOmics DIABLO package (green lines; correlation coefficient = 0.7, Bonferroni-adjusted p<0.05). (**b**) Abundance of chaperone protein AELOFPMA_23737 and (**c**) urease subunit alpha GFDOOBDM_36378 were increased by 12 weeks CS-exposure. (**d**) Abundance of oxidative stress-responsive proteins GDJJNAEL_42303 and (**e**) BGMILIAL_35735 demonstrated increases and decreases after CS-exposure respectively, which was alleviated by FMT. (**f**) Abundance of energy metabolism proteins AAEHFLO_09769 and (**g**) EAAGGHIE_49748 were increased by CS-exposure, which was alleviated and enhanced by FMT respectively. **h-l,** Mice were exposed to CS or air for 12 weeks, and metabolites in cecum contents assessed. Components of the glycolysis and gluconeogenesis, or starch and sucrose metabolism pathways were downregulated in CS-exposed mice. N = 3 (**a-g**) or 8 per group (**h-l**). Data presented as mean +/- standard error of the mean. * = p<0.05; ** = p<0.01; *** = p<0.001; **** = p<0.0001 using one-way ANOVA with Holm-Sidak’s post-hoc analysis (**c-g**) or unpaired Student’s T test on log-transformed data (**h-l**).

Of these proteins correlated with bacterial genera, 12 were significantly altered in at least one experimental group (Table S15). Chaperone protein AELOFPMA_23737 and urease subunit alpha GFDOOBDM_36378 were increased by 12 weeks CS-exposure, but not in smoking cessation, and were not alleviated by FMT (Fig. 5b-c).

Four proteins were altered by both CS-exposure and FMT (Fig. 5d-g). Alkyl hydroperoxide reductase subunit C GDJJNAEL_42303, an antioxidant protein, was increased after 12 weeks of CS-exposure and alleviated by smoking cessation and FMT (p=0.052) (Fig. 5d). Rubrerythrin BGMILIAL_35735, also implicated in oxidative stress responses, had a reverse pattern and was decreased after 12 weeks of CS-exposure and after smoking cessation, and increased by FMT (Fig. 5e). Phosphoglycerate kinase AAEHAFLO_09769 and malate dehydrogenase EAAGGHIE_49748, implicated in glucose metabolism and energy generation, were both increased by 12 weeks of CS-exposure and, to a lesser extent, after smoking cessation (Fig. 5f-g). While FMT alleviated the CS-induced increase in AAEHAFLO_09769, it further increased EAAGGHIE_49748. Metabolite profiling in the cecum contents of a separate cohort of CS-exposed mice identified that glycolysis, gluconeogenesis, starch and sucrose metabolism components were downregulated (Fig. 5h-l). Thus, proteomics and metabolomics implicated a downregulation of glucose and starch metabolism in CS-exposed mice, which was altered by FMT.

### Dietary resistant starch alleviated inflammation and emphysema in experimental COPD

Given the changes in microbial glucose and starch metabolism, we assessed whether dietary supplementation with complex carbohydrates improved disease outcomes. Mice were exposed to CS (8 weeks), and fed either control diet or an equivalent diet with all carbohydrates as resistant starch (high amylose maize starch) which is not digested by the host, increasing the availability to microbiota.^33^ Resistant starch alleviated CS-induced airway inflammation and prevented CS-induced increases in alveolar diameter (Fig. 6a-e), consistent with the protective effects of FMT.

**Figure 6:**
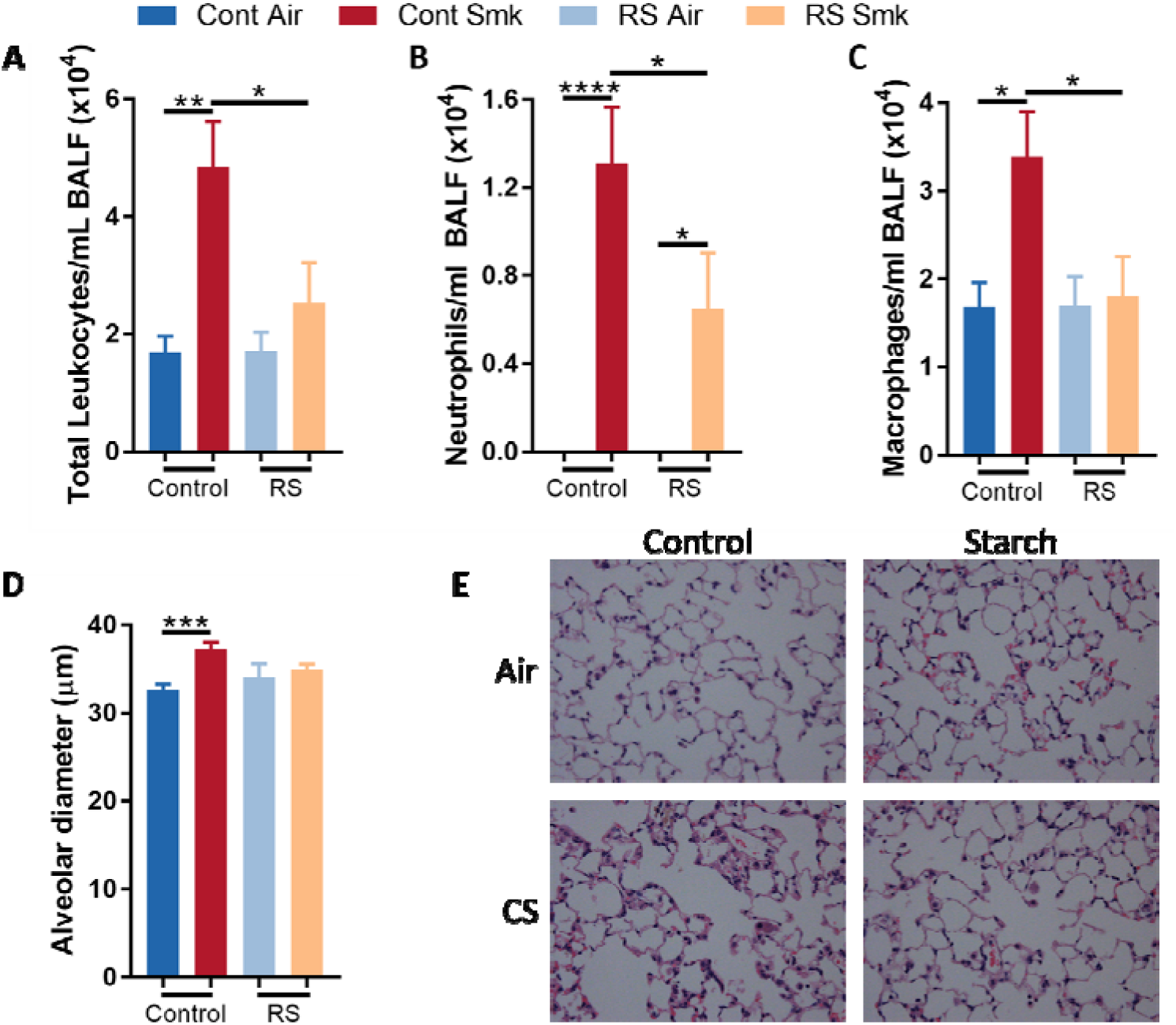
High resistant starch intake alleviated cigarette smoke (CS)-induced inflammation and emphysema. **a-d,** Mice were exposed to CS or normal air for 8 weeks. Beginning 2 weeks prior to exposure, mice were fed a control or resistant starch (RS) diet. RS diet alleviated CS-induced increases in (**a**) total leukocytes, (**b**) neutrophils and (**c**) macrophage in bronchoalveolar lavage fluid (BALF), and prevented (**d, e**) CS-induced alveolar destruction. N = 6-14 per group. Data presented as mean +/- standard error of the mean. * = p<0.05; ** = p<0.01; *** = p<0.001; **** = p<0.0001 using one-way ANOVA with Holm-Sidak’s post-hoc analysis.

### Inulin supplementation reduced symptoms and exacerbations in human COPD

To explore whether these findings translated into humans, we performed a small, preliminary pilot study with a randomized, double-blind, placebo-controlled study in COPD patients receiving supplements of inulin, a common fermentable fiber, for four weeks. There were no differences between placebo-(N=7) and inulin-treated (N=9) patients in age, gender, BMI, smoking history, lung function, proportion of frequent exacerbators (≥2 exacerbations in 12 months), medication use or seasonality of intervention (Table 1). Oral inulin resulted in fewer self-reported exacerbations (worsening or respiratory symptoms requiring additional pharmaceutical intervention) during the intervention period (1/9 patients, 11%) than placebo (4/7 patients, 57%) (Table 2) and improved health-related quality of life measured by two validated instruments; the COPD Assessment Test (CATest) and the St Georges Respiratory Questionnaire (SGRQ). Both CATest and SGRQ scores were lower (improved symptoms) after inulin treatment than at baseline, exceeding the Minimally Important Clinical Difference.^34^ There was no reduction in either score after placebo (Table 2), but only the change in CATest score was significantly different in inulin-treated patients compared to placebo.

**Table 2:** Exacerbation frequency, COPD assessment test (CAT) and St Georges Respiratory Questionnaire symptom scores (SGRQ) in COPD patients receiving 4 weeks placebo (N=7) or inulin (N=9) supplement.

|  | Placebo |  | Inulin |  | P-value |
| --- | --- | --- | --- | --- | --- |
|  | Mean | SD | Mean | SD |  |
| Number of exacerbations | 4 (57%) | N/A | 1 (11%) | N/A | **\*0.048** |
| Pre-treatment CATest score | 14.33 | 1.16 | 17.13 | 7.18 | >0.99 |
| Post-treatment CATest score | 16.00 | 1.00 | 12.88 | 5.33 | >0.99 |
| Δ CATest score | 1.67 | 2.08 | -4.25 | 2.49 | **\*0.01** |
| Pre-treatment SGRQ score | 40.49 | 13.08 | 42.52 | 16.94 | >0.99 |
| Post-treatment SGRQ score | 41.91 | 3.24 | 36.17 | 14.51 | >0.99 |
| Δ SGRQ total score | 1.43 | 9.94 | -6.34 | 6.07 | **0.27** |

## Discussion

CS-induced changes were broadly similar across several experiments, with *Muribaculaceae Desulfovibrionaceae* and *Lachnospiraceae* members increased after 8 and 12 weeks and correlating with disease phenotypes. FMT only partially alleviated CS-induced changes in microbiome composition, likely due to continued selective pressure with continuous smoke exposure or incomplete microbiota replacement using the bedding swap FMT. Bedding swaps, while not the most controlled method of FMT,^35^ were used in most experiments due as repeated oral gavage affects histopathology and lung function in mice.^36^ Nevertheless, FMT reproducibly alleviated inflammation, emphysema, impaired lung function, and gastrointestinal pathology and our initial findings were validated by FMT using oral gavage, indicating that gut microbiota can have substantial effects on disease. Minor differences in microbiome changes between studies were expected due to natural variability,^37^ and cage/batch effects were controlled by microbiome normalization. Together, these findings demonstrate effective alleviation of disease by FMT and associations with bacterial taxa across multiple independent experiments. Reproducibility between experimental models of COPD has proven challenging,^10, 11^ as has the correspondence between changes observed in murine models and samples from human COPD pateints. COPD patients have reduced abundance of *Desulfovibrionaceae* member *Desulfovibrio piger*, but increased abundance of select *Lachnospiraceae* species including *Sellimonas intestinalis* that negatively correlated with lung function,^8^ while CS-exposure in mice increased *Lachnospiraceae* abundance.^11^ This is the first time that *Muribaculaceae* species have been implicated in COPD patients or animal models. Although more predominant in the murine microbiome, this family is present in humans and may have roles in disease.^8^ Two *Muribaculaceae* species, *M. intestinale* and *UBA7173 sp001689685*, negatively correlated with inflammation and emphysema, highlighting the importance of assessing the microbiome at higher taxonomic resolution.^10^

Of the CS-associated species, *Amulumruptor sp001689515* and *Mailhella sp003512875* positively correlated with inflammation and emphysema. Gram-negative *Mailhella sp003512875* also positively correlated with colon *Tlr4* expression. Fecal filtrates from CS-exposed mice impaired TLR4 agonist-induced inflammatory responses *in vitro*, suggesting that microbial stimulation of TLRs may influence the gut-lung axis and/or facilitate the persistence of pathogenic taxa. We previously reported opposing roles for lung TLR4 (pathogenic) and TLR2 (protective) in COPD^18^ and here we provide evidence that their signaling in the colon impacts pathogenesis, consistent with the protective effects of *P. goldsteinii* mediated by reduced activation of TLR4.^11^ Moreover, changes in TLR3, which recognizes damage associated-molecular patterns, suggests an association with colonic pathology.^38^ Whether changes in colonic TLR expression are a result of altered gene expression in resident cells or infiltration of high TLR-expressing lymphocytes is not clear.

FMT did not alleviate CS-induced changes in bone marrow leukocytes but did alleviate increases in blood Ly6C^lo^ (non-classical) monocytes and depletion of blood B cells and splenic CD8^+^ cDCs. Non-classical monocytes, associated with tissue repair or autoimmune damage,^39^ are increased in severe COPD^40^ and thus the FMT-mediated reduction is likely associated with suppressed alveolar destruction. CD8^+^ cDCs, particularly in the spleen, are implicated in immune tolerance, phagocytosis and cross-presenting antigens, suggesting an association with increased blood B cells in FMT-treated mice.^41^ Splenic cDC numbers are increased in models of acute lung injury,^42^ demonstrating a spleen-lung axis regulated by gut microbiota.

In the fecal proteome, increased chaperone protein DnaK (AELOFPMA_23737) and urease subunit alpha (GFDOOBDM_36378) after 12 weeks CS may be in response to heavy metals which rapidly reduce after smoking cessation.^43–45^ Hydrolysis of urea by urease increases nitrogen availability which has been linked to the persistence of deleterious bacteria and inflammation.^45^

Alkyl hydroperoxide reductase GDJJNAEL_42303 and rubrerythrin BGMILIAL_35735 deactivate hydrogen peroxide to permit bacterial survival.^46, 47^ Increased alkyl hydroperoxide reductase in response to CS-induced oxidative stress^33^ may explain the expansion of *Desulfovibrionaceae* that thrive in oxidative conditions,^48^ and antioxidants are postulated to maintain the mucous barrier in COPD patients.^33^ That rubrerythrin was reduced by CS and increased by FMT indicates that the local environment influences oxidative stress responses and microbial adaptability. For example, rubrerythrin uses iron as a co-factor^46^ whereas alkyl hydroperoxide reductase is repressed in high iron environments.^47^ Many COPD patients have iron deficiency^33^ and lower hepcidin levels in CS-exposed mice increase absorption and reduce iron in the intestinal lumen,^49^ and may contribute to the different oxidative stress responses and expansion of pathogenic taxa in COPD.

Phosphoglycerate kinase AAEHAFLO_09769 and malate dehydrogenase EAAGGHIE_49748 are involved in glycolysis and gluconeogenesis^50, 51^ and their abundance is increased in nutrient restriction, including by *Lachnospiraceae* spp..^50, 51^ Nicotine is an appetite suppressant^52^ and reduced food consumption alters intestinal microbiota and predisposes to *Streptococcus pneumoniae* infection,^33^ a major cause of COPD exacerbations and lung dysbiosis.^5^

Metabolomics also implicated impaired carbohydrate metabolism. Complex carbohydrates like resistant starch and inulin are poorly digested by the host, improving availability to microbiota stimulating anti-inflammatory SCFAs.^6, 33^ High fiber intake is associated with lower risk of developing COPD,^33^ and inulin reduced airway inflammation in asthma.^53^ Resistant starch reduced inflammation and emphysema in experimental COPD, and our pilot study in COPD patients indicated that even a short period of inulin supplementation may reduce exacerbations and symptoms. However, examination of exacerbation frequency during the short period of intervention and assessment (4 weeks) should be interpreted with caution. Trials in larger cohorts of COPD patients, and over longer periods of time, are needed to definitively demonstrate alleviation of symptoms and exacerbations and to progress these findings into clinical practice. Although inulin differed from the resistant starch used in the mouse model, its wide use as an established prebiotic in humans improves translatability for these future studies.^53^

Jang *et al*. showed limited evidence for FMT alleviating whole-body CS-exposure/poly I:C-induced emphysema which were associated, but not correlated, with increased *Lactobacillaceae* and decreased *Bacteroidaceae* identified through 16S rRNA gene sequencing.^10^ Whole body CS-exposure coats the fur and grooming will unnaturally affect gut microbiota.^54^ Without appropriate controls (no CS or poly I:C alone) and baseline microbiome normalization or sampling, it was not possible to discern the different effects of CS and poly I:C, a TLR3 agonist, or determine whether changes in microbiota were due to confounding factors including cage effects,^37^ accelerated ageing in COPD and the maturation of the microbiota,^1^ or the effects of PBS/saline vehicle gavage on gut microbiota, immune responses and lung histopathology.^36^ That study was not translatable, as it did not assess the most clinically relevant feature of COPD (impaired lung function) or assess efficacy in humans.^10^ The lack of human translation has been a limitation of other models of experimental COPD^11^, particularly given that experiments rarely consider the impacts in the context of smoking cessation, the current first-line treatment for COPD.

Our nose-only CS-exposures accurately reflects human exposures and disease mechanisms,^4^ and included microbiome normalization and baseline microbiome assessment and demonstration of therapeutic benefits using FMT by both bedding swap and oral gavage to discriminate between disease-associated changes and those from confounding factors. Higher resolution metagenomic sequencing and correlation of individual taxa with disease features allowed species-level classification to elucidate the different, and sometimes contrasting, effects of family members such as CS-associated *Amulumruptor sp001689515* and air/FMT-associated *M. intestinale* and *UBA7173 sp001689685* from the *Muribaculaceae* family. The efficacy of FMT was demonstrated in several contexts, including mild and severe disease, gastrointestinal co-morbidities and providing additional benefits to smoking cessation by alleviating persistent increases in macrophages that drive progressive disease.^12^ Translatability was enhanced by assessing lung function and extending our work in a pilot study to demonstrate clinically relevant alleviation of symptoms in patients. Overall, our study provides strong evidence for the therapeutic efficacy of FMT and complex carbohydrate supplementation which could be acceptable interventions for COPD patients, that may become more effective when combined with smoking cessation.^55^

## Conclusions

We demonstrate that dysbiosis, including increased *Muribaculaceae*, *Desulfovibrionaceae* and *Lachnospiraceae* species, contributes to disease pathogenesis and FMT alleviates hallmark features of CS-induced COPD. These changes may be affected by microbial energy metabolism, and complex carbohydrate supplementation protects against experimental disease and reduced exacerbations and symptoms in humans. This work opens new avenues towards effective microbiome-targeting therapies in COPD.

## ACKNOWLEDGEMENTS

**Acknowledgments and funding:** We thank the Rainbow Foundation and Felicity and Michael Thomson for their support, Kristy Wheeldon, Emma Broadfield, Bradley Mitchell and Nathalie Kiaos for their help with animal models, and Nathan Smith for his help with proteomics analysis. This work and PMH were supported by grants and fellowships from the National Health and Medical Research Council (NHMRC) of Australia (1059238, 1079187, 1175134), the Rainbow Foundation, Australian Research Council (110101107), Cancer Council of NSW, University of Newcastle and University of Technology Sydney and The Prince Charles Hospital Foundation (RF2017-05, INN2018-30). J.L.S. was supported by German Research Foundation (DFG) under Germany’s Excellence Strategy (EXC2151-390873048) as well as under SFB 1454 (432325352), the BMBF-funded excellence project Diet–Body–Brain (DietBB), EU grant under Horizon 2020 DiscovAir (874656), and the EU project SYSCID (733100).

