## Supplementary files for "Fecal microbial transfer and complex carbohydrates mediate protection against COPD"

**Supplementary Figures:**

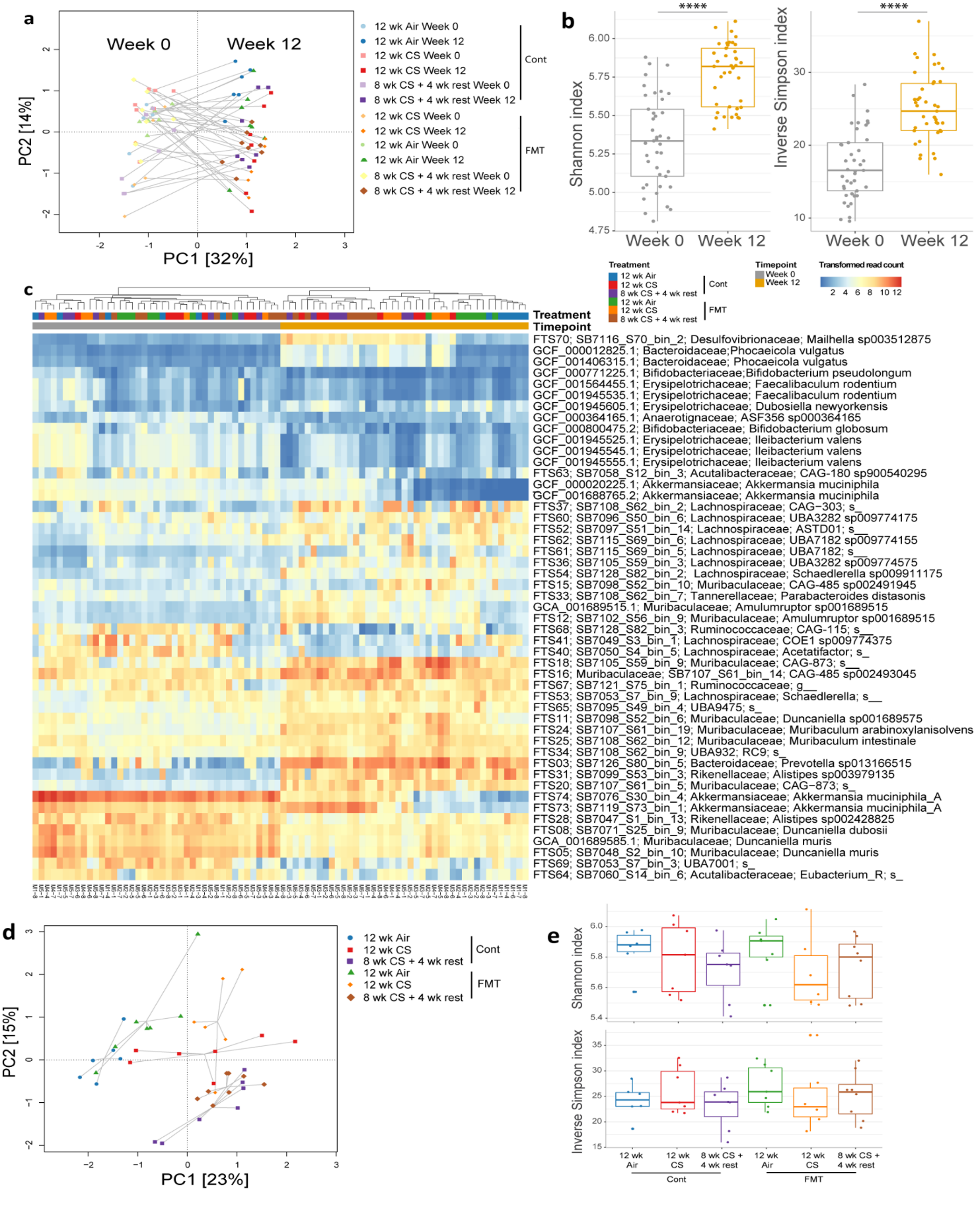
**Figure S1. Fecal microbiome profiles are significantly altered across time with cigarette smoke (CS)-exposure and fecal microbiota treatment (FMT).** **a-e,** Mice were exposed to CS (12 wk CS) or normal air (Air) for 12 weeks, or 8 weeks CS followed by 4 weeks with normal air (8 wk CS+ 4 wk rest). Mice also received FMT through transfer of soiled bedding twice per week or were maintained in their own bedding (Control) for 12 weeks. Fecal samples from baseline (week 0) and at the end of the experiment (week 12) were analyzed using shotgun metagenomics. (**a**) Principal component analysis of all mice demonstrated shifts in microbiome composition with time, smoke-exposure and FMT based on genome level read mapping counts, log-cumulative-sum-scaling (CSS) transformed. (**b**) Simpson and Shannon diversity indices for all mice demonstrated increasing diversity over time based on mapping counts across all genomes transformed using the DESeq2 scaling factor. (**c**) Heatmap displaying significantly different genomes (Benjamini-Hochberg adjusted P<0.001, log2 fold change >1.5) between week 0 and week 12 as log-CSS transformed read counts. (**d**) Principal component analysis of week 12 mice demonstrated significant separation between groups using PERMANOVA of Bray-Curtis distances. (**e**) Simpson and Shannon diversity indices indicating no differences in diversity between experimental groups at week 12. N = 8 per group. **** = p<0.0001 using a Wilcoxon rank sum test.

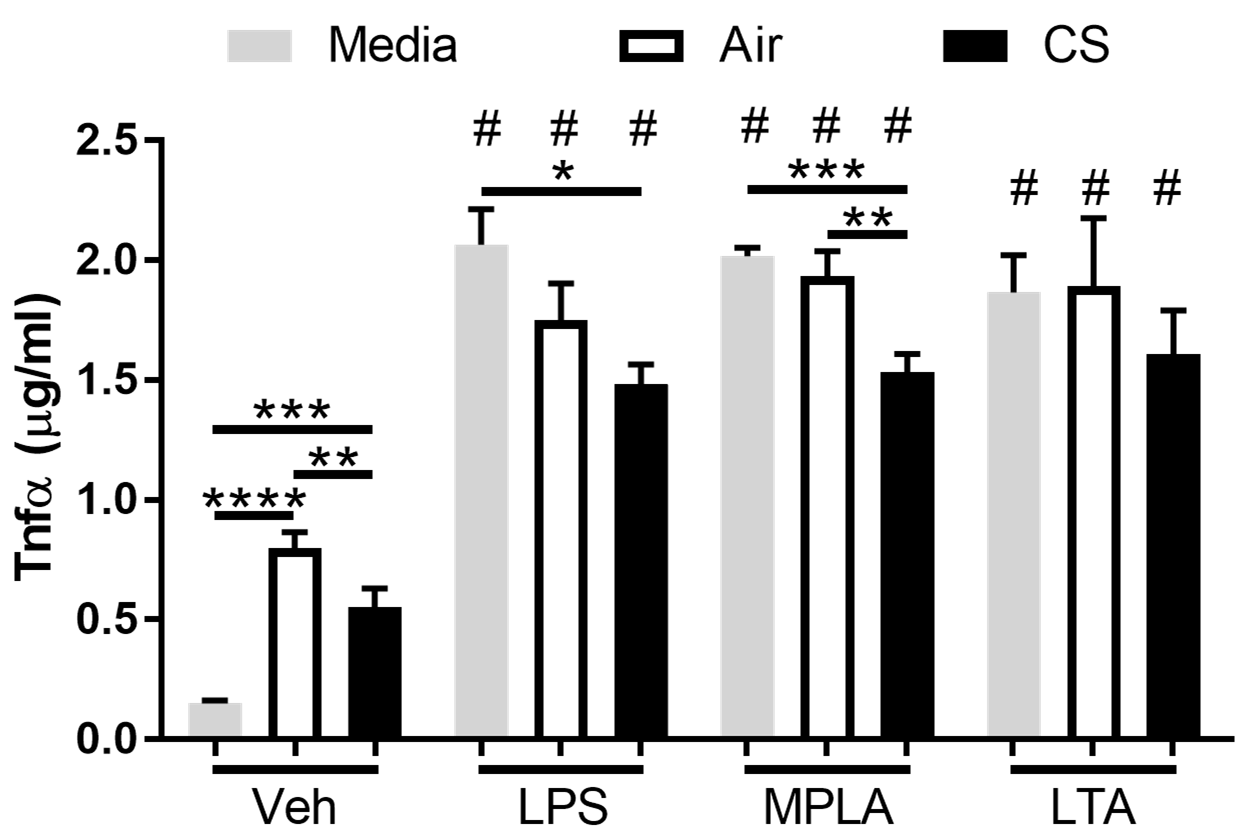

**Figure S2: Cigarette smoke (CS)-induced dysbiosis impaired TLR4 agonist-induced TNFα production in monocytes.** Raw 264.7 monocytes were incubated for 24 hours with sterile-filtered faecal homogenates from mice exposed to CS (black) or normal air (white) for 12 weeks, or were maintained in media (grey). In the last 4 hours of incubation, cells were incubated with lipopolysaccharide (LPS), monophosphoryl lipid A (MPLA), lipoteichoic acid (LTA) or vehicle (Veh; 0.5% dimethylsulfoxide). Incubation with faecal homogenates induced TNFα production, which was lower with faeces from CS-exposed mice. LPS- and MPLA-induced TNFα production was also lower in cells incubated with faeces from CS-exposed mice. N = 3-9 per group. Data presented as mean +/- standard error of the mean. ** = p<0.01; *** = p<0.001; **** = p<0.0001; # = p<0.0001 *vs.* Veh-treated cells using one-way ANOVA with Holm-Sidak’s post-hoc analysis.

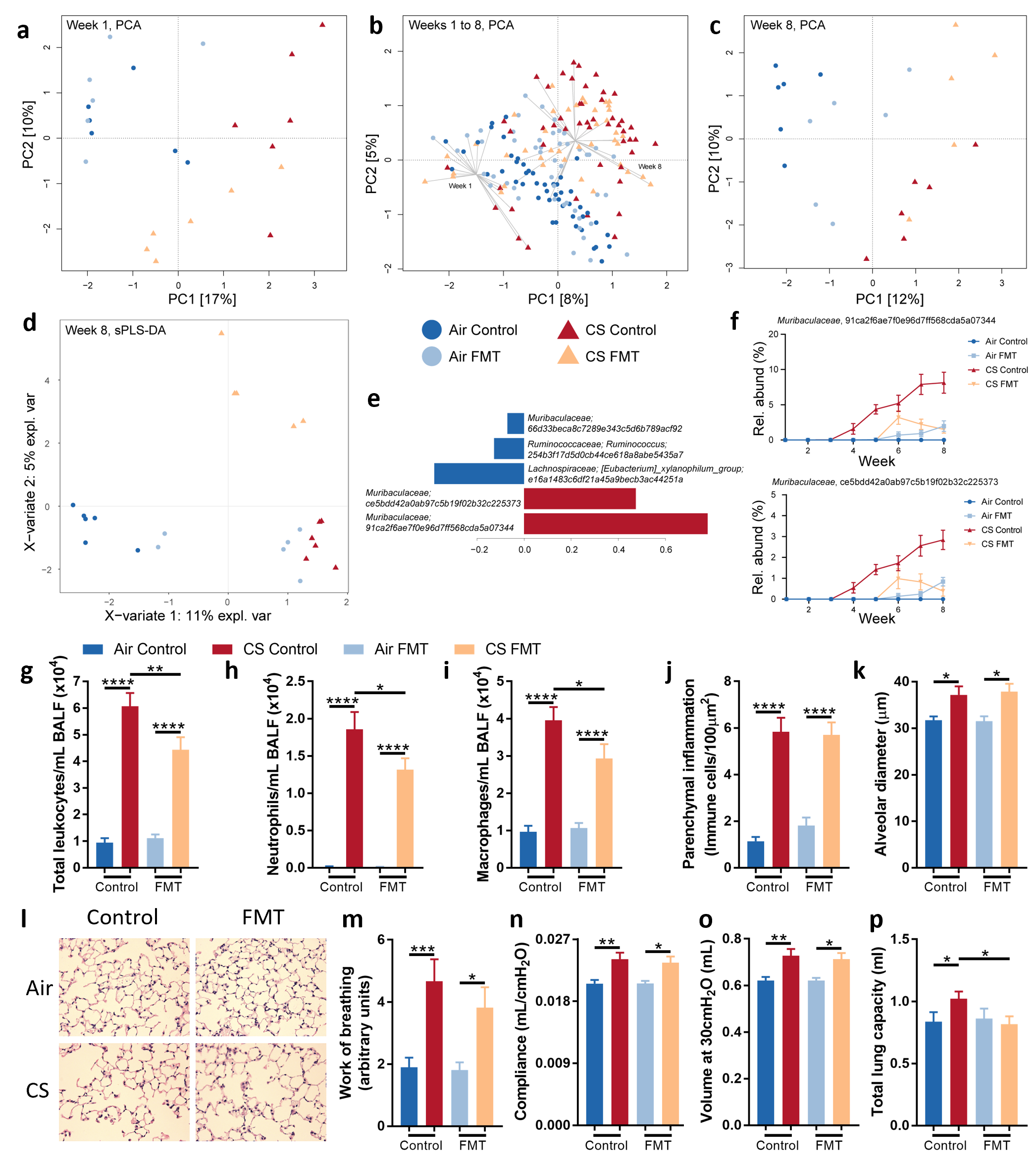

**Figure S3.** **Cigarette smoke (CS)-exposure was associated with a shift in the faecal microbiome, and faecal microbiota transfer (FMT) suppressed the development of experimental COPD**. Mice were exposed to CS or normal air for eight weeks and received FMT through transfer of soiled bedding or were maintained in their own bedding (Control). **a-f,** Faecal samples were collected weekly and analyzed by 16S rRNA amplicon sequencing. (**a**) Principal component analysis of mice following one week of CS-exposure and/or FMT, across (**b**) all eight weeks of the experiment and at (**c**) week eight following completion of the experiment demonstrated a clear distinction in microbiota composition of the different experimental groups. (**d**) Multivariate analysis (sPLS-DA) at week eight demonstrated distinction between air- and CS-exposed groups along component 1, (**e**) with several species contributing to this separation. (**f**) Relative abundance of *Muribaculaceae* sequence variants increased in CS-exposed mice but was alleviated by FMT, contributing to separation of sPLS-DA across the eight-week experiment. **g-p,** Hallmark features of COPD assessed at week eight. CS-exposure increased (**g-i**) total leukocytes, neutrophils and macrophages in bronchoalveolar lavage fluid (BALF), (**j**) immune cells in parenchyma, (**k-l**) alveolar diameter, and (**m-p**) lung function parameters of work of breathing, compliance, volume and total lung capacity. FMT alleviated CS-induced increases in BALF total leukocyte, neutrophil and macrophage numbers and total lung capacity. N = 5-6 per group (**a-f**) or 14-18 per group from 3 independent experiments (**g-p**). Data presented as mean +/- standard error of the mean. * = p<0.05; ** = p<0.01; *** = p<0.001; **** = p<0.0001 using one-way ANOVA with Holm-Sidak’s post-hoc analysis (**g-p**).

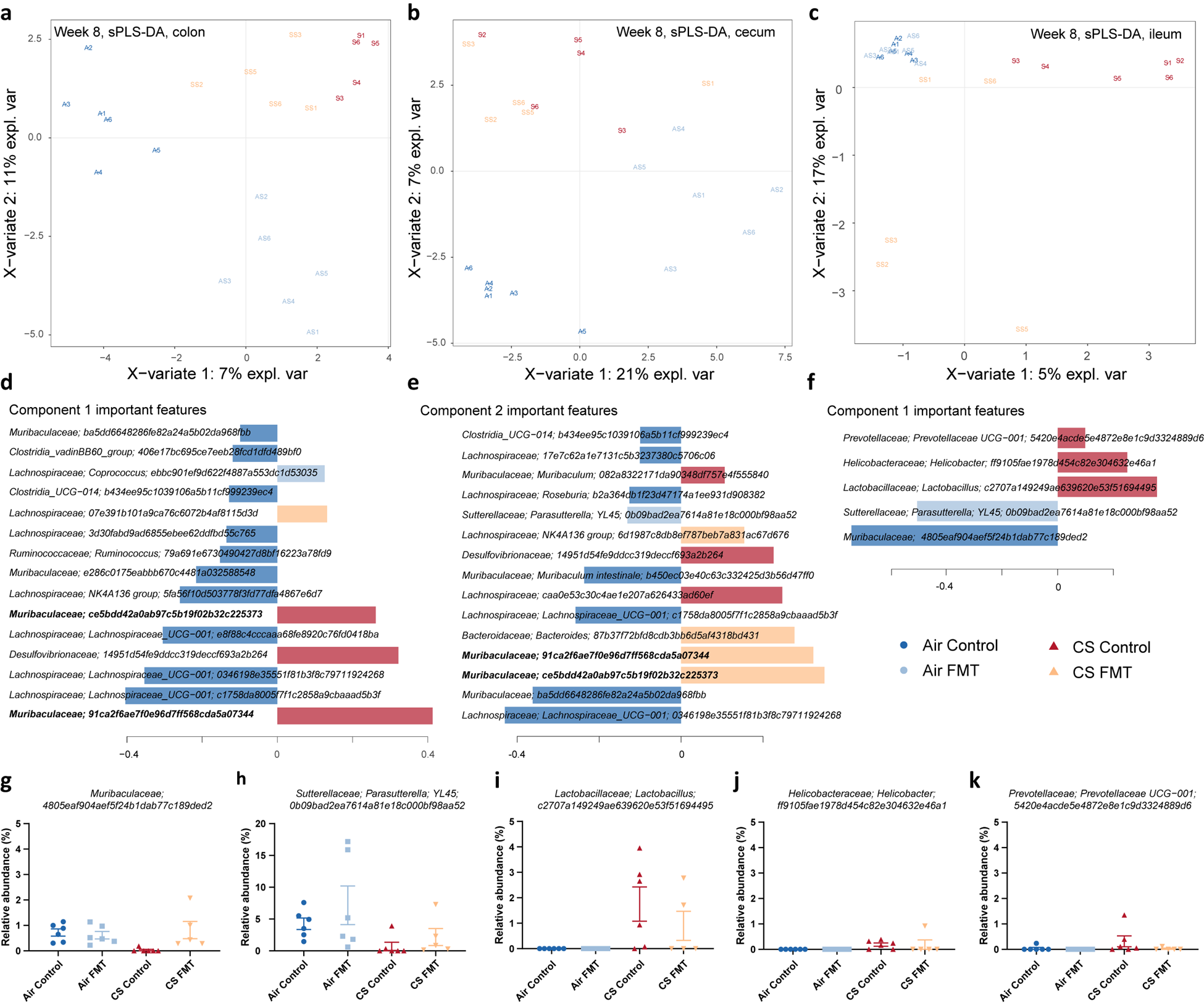

**Figure S4. Analysis of ileum samples reveals distinction between cigarette smoke (CS)-exposed groups based on faecal microbiota transfer (FMT) treatment.** **a-k,** Mice were exposed to CS or normal air (Air) for eight weeks, and received FMT through transfer of soiled bedding or were maintained in their own bedding (Control). Multivariate analysis (sPLS-DA) at week eight based on 16S rRNA amplicon sequencing indicated CS-induced separation of microbiome samples, resembling that observed in faecal samples, in the (**a**) colon and (**b**) caecum but not the (**c**) ileum, although the latter did demonstrate a difference between CS control and CS FMT groups. (**d**) Species contributing to separation along the CS-associated component 1 of **a** and (**e**) component 2 of **b**. (**f**) Species contributing to separation along the air-associated component 1 of **c**. (**g-k**) Relative abundance of air-associated ASVs from the ileum from CS FMT mice trended towards the levels of air-exposed control mice. N = 5-6 per group.

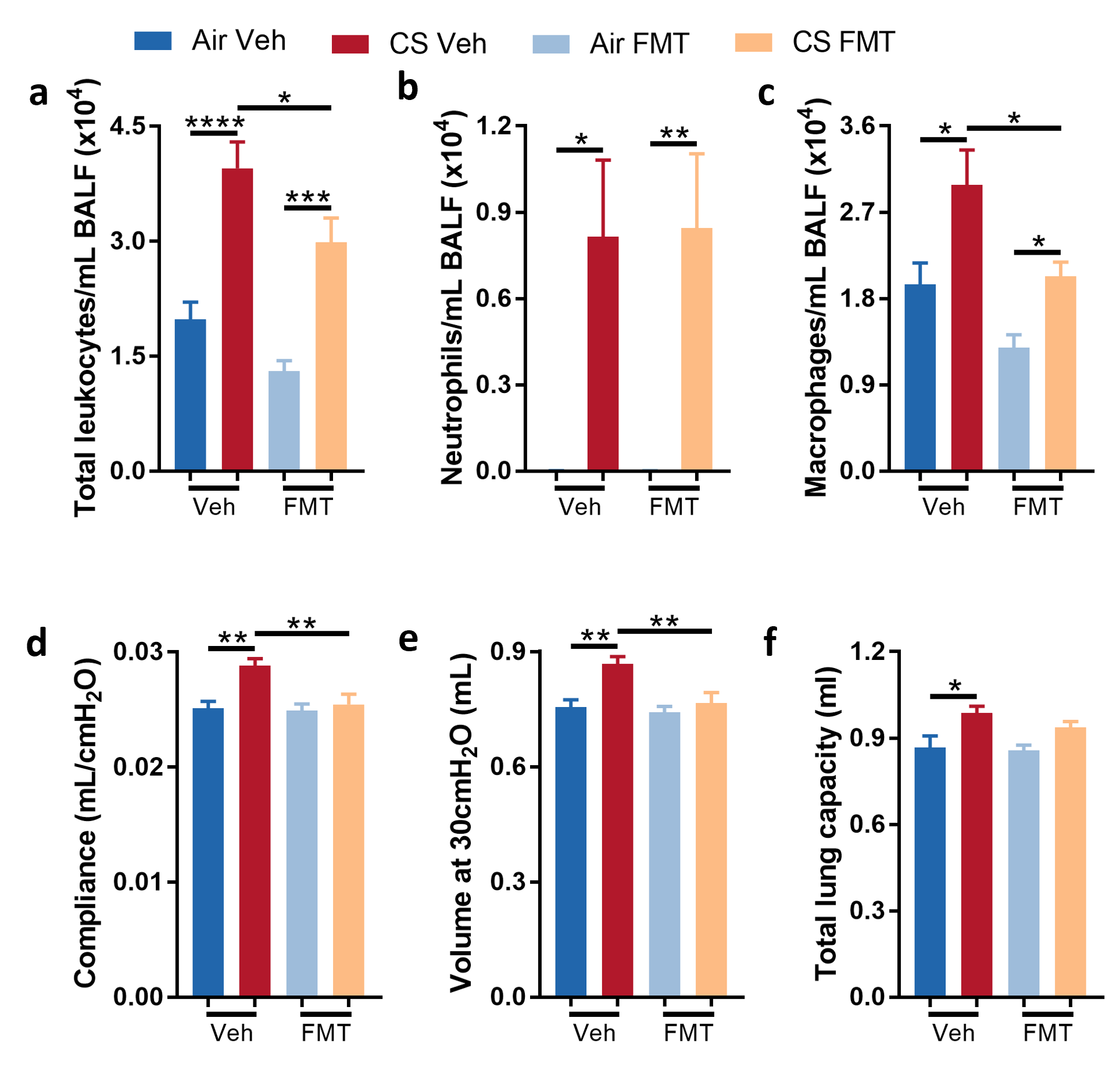

**Figure S5. Faecal microbiota transfer (FMT) by oral gavage suppressed the development of experimental COPD**. **a-f,** Mice were exposed to cigarette smoke or normal air for eight weeks and received FMT or vehicle (Veh; phosphate buffered saline + 0.05% L-cysteine) through oral gavage. (**a-c**) CS-exposure increased total leukocytes, neutrophils and macrophages numbers in bronchoalveolar lavage fluid (BALF), and (**d-f**) the lung function parameters compliance, volume and total lung capacity. FMT alleviated CS-induced increases in BALF total leukocytes and macrophages, and lung compliance and volume. N = 10-12 per group from 2 independent experiments. Data presented as mean +/- standard error of the mean. * = p<0.05; ** = p<0.01; *** = p<0.001; **** = p<0.0001 using one-way ANOVA with Holm-Sidak’s post-hoc analysis.

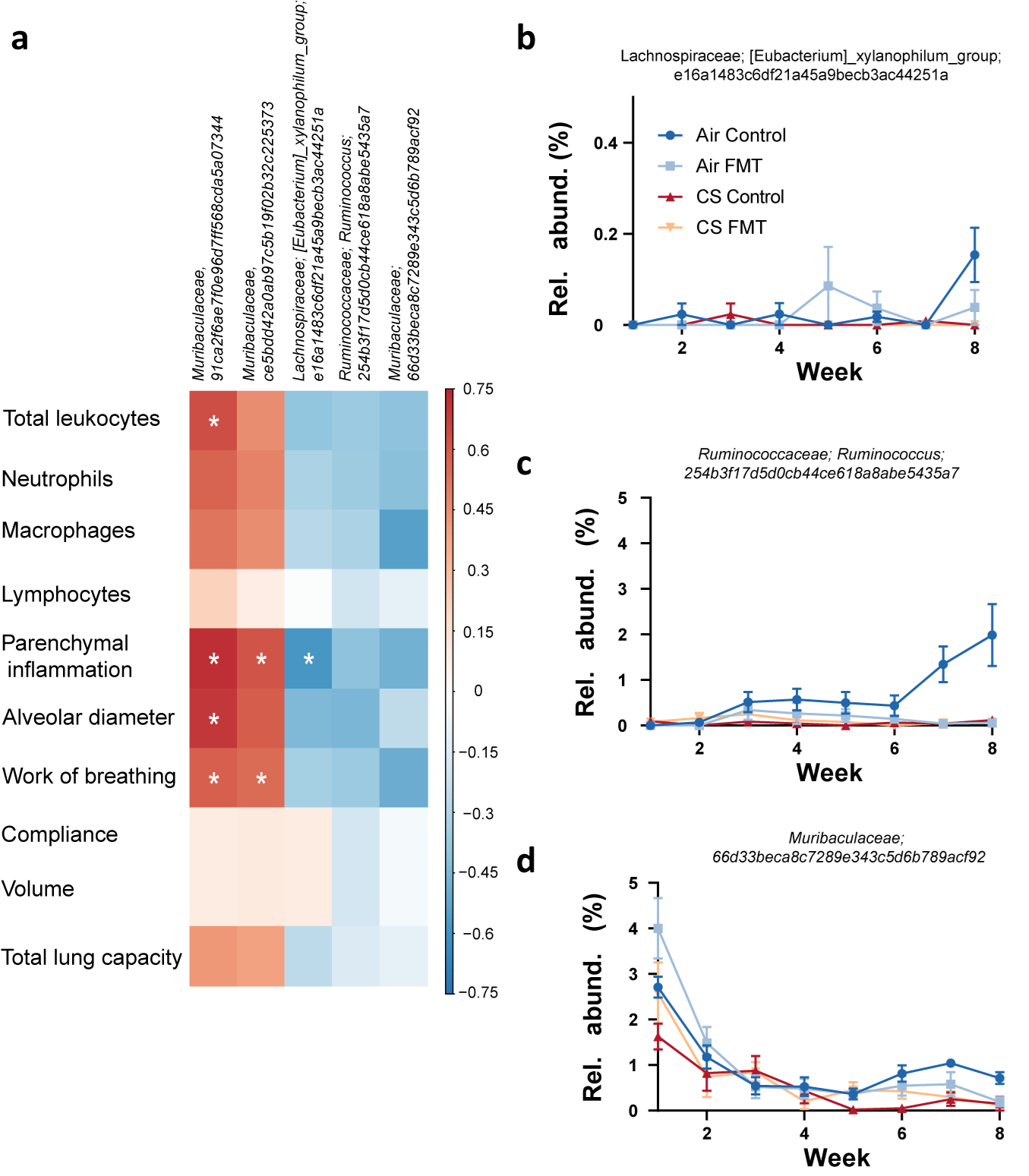

**Figure S6: A cigarette smoke (CS)-associated *Muribaculaceae* amplicon sequence variant (ASV) correlates with a subset of phenotypic measures.** **a-d,** Mice were exposed to CS or normal air (Air) for eight weeks and received faecal microbiota transfer (FMT) through transfer of soiled bedding or were maintained in their own bedding (Control). Faecal samples were collected weekly and analyzed by 16S rRNA amplicon sequencing. (**a**) Spearman’s rho calculated between centered log-ratio transformed ASV relative abundance and phenotypic scores indicated correlation between *Muribaculaceae 91ca.* and a subset of phenotypic measures. (**b-d**) Relative abundances of air-associated ASVs from Fig. S2e across the eight-week experiment were variable. N = 5-6 per group. * = p<0.05 using Spearman’s correlation.

**Supplementary Methods:**

**Mice, CS-exposure, microbiome transfer and diet studies**

Female C57BL/6 mice (3-5 weeks old) were obtained from the Animal Services Unit, The University of Newcastle (Newcastle, Australia). Upon arriving in the facility, the baseline microbiome was normalized for three weeks by pooling and mixing all soiled bedding and redistributing to all cages (twice weekly) and co-housing mice from different experimental groups weekly. Microbiome normalization was not performed for validation experiments (8 weeks CS with FMT).

Mice were exposed to normal room air or CS from twelve 3R4F reference cigarettes (University of Kentucky, Lexington, KY) in a custom-designed, nose-only inhalation apparatus, twice per day, 5 days per week, for 8 or 12 weeks, as previously described.^12-18, 44, 45^ In some longer experiments, mice exposed to CS for 8 weeks were rested for the final 4 weeks of the experiment to model smoking cessation. Twice per week, soiled bedding from air-exposed mice was mixed with clean bedding (1:2 ratio) and transferred to the cages of smoke-exposed mice. Conversely, the soiled bedding from 12-week CS-exposed mice was mixed with clean bedding and transferred to the cages of air-exposed mice. Control groups (without FMT) received their own soiled bedding mixed with clean bedding. Feces were collected weekly, with at least 3 fresh fecal pellets collected and stored at -80°C until processing.

For FMT by oral gavage, fresh feces were collected from normal air-exposed mice and placed immediately in phosphate buffered saline (PBS) with 0.05% L-cysteine to preserve anaerobic bacteria (100mg/1.5mL). Feces was gently homogenized by hand and debris removed by centrifugation (800xg, 3 min). Resulting supernatants were administered by oral gavage, beginning in the first week of smoke exposure. For diet studies, mice were fed a conventional semi-pure diet (AIN93G) or a resistant starch diet (SF11-025; Specialty Feeds, WA, Australia) *ad libitum* commencing 2 week prior to CS-exposure and maintained until the end of experiment. All experiments were approved by University of Newcastle Animal Ethics Committee.

**DNA extraction and metagenome assembly and binning**

DNA was extracted from 50-100 mg of fecal material using an initial bead beating step and extraction using a Maxwell 16 Research Instrument and Tissue DNA Kit (Promega, USA) according to the manufacturer’s protocol. DNA concentrations were measured using a Qubit assay (Life Technologies, USA) and were adjusted to a concentration of 5 ng/ul. Library preparation was performed using Nextera DNA Library Preparation Kits (Illumina, USA) and samples sequenced across two runs on an Illumina NextSeq500 instrument generating an average of 3 Gbp of 150 bp paired-end reads per sample. Each sample was included on both sequencing runs and data combined. Contaminating host reads were removed by mapping against the mouse genome (*Mus musculus* GRCm38.p5) using BWA v0.7.12^1^ requiring a minimum alignment length of 30 bases and maximum of 15 clipped bases for reads to be considered of mouse origin. Raw host-free reads from each sample were assembled using Spades v3.12.0^2^ with the --meta flag. Reads were mapped to each resulting assembly using BamM v1.7.3 (https://github.com/ecogenomics/BamM) and bins produced using Metabat v2.12.1,^3^ MaxBin^4^ and GroopM2 v2.0.0-1^5^ implemented within the ensemble binning tool UniteM v0.0.15 (<https://github.com/dparks1134/UniteM>). Contamination and completeness of bins from all samples were assessed using CheckM v1.0.11.^6^ Bins with completeness >80% and contamination <7% were retained and de-replicated using dRep v2.05^7^ with default settings (99% identity), skipping quality filtering, to produce the final set of metagenome-assembled genomes (MAG). Taxonomy was determined using GTDB-Tk v0.3.0 and v1.5.0^8^ with GTDB releases 04-RS89 and 06-RS202.^9^

**Metagenomic community profiling**

Reads for each sample were mapped to a de-replicated set of 18,342 genomes from NCBI (GTDB release 03-RS83) using BamM with minimum seed length of 25. Genomes with >1x coverage of >1% of the genome and overall coverage of 0.01X, as determined using Mosdepth v0.2.3,^10^ were retained and combined with a de-replicated set of recovered MAGs for assessment of community composition. Read counts for the final genome set was determined for each sample *via* mapping using BamM with a minimum seed length of 25 bases and filtering for minimum mapping percentage identity of 95%. Per genome read counts were scaled to account for genome size whilst maintaining the raw unmapped read percentage for each sample as a reflection of unrepresented diversity. Relative abundance was calculated using scaled read counts as a fraction of total non-host reads per sample. α-diversity was calculated using QIIME v1.8.0^11^ with read counts per-genome normalised to account for library size using DESeq2 v1.20.0.^12^

**Statistical analysis of metagenomic data**

Principal component analysis was performed using the R package vegan v2.5-1^13^ on data transformed using log cumulative-sum-scaling (log-CSS) implemented within metagenomeSeq v1.22.0.^14^ Differential abundance between sample groups was determined using DESeq2 using read counts per genome scaled to account for genome size and Benjamini-Hochberg adjustment for multiple comparisons. sPLS-DA analysis was conducted using the R package mixOmics v6.3.2^15^ with centered log-ratio transformed relative abundance (adding pseudo count one order of magnitude below the lowest non-zero value values) with 50x4-fold cross-validation. Host phenotypes were tested for association with microbiome composition using the envfit function based on NMDS ordination of Bray-Curtis distances calculated using the metaMDS function, both implemented within the vegan R package. Significance was determined based on 10,001 permutations and resulting P values were adjusted for multiple comparisons using the Benjamini-Hochberg method. Samples with missing scores were removed. Spearman’s rho was calculated using the ‘corr.test’ function within the R package psych v1.8.12^16^ based on centered log ratio transformed genome relative abundance and raw phenotypic data. Correlation matrix was produced using the ‘corrplot’ function with the R package corrplot v0.84.^17^

**Assessment of experimental COPD: airway inflammation, emphysema-like alveolar enlargement**

Airway inflammation was quantified by total and differential enumeration of immune cells in BALF as previously described.^18-20^ Briefly, two 0.4 ml washes with PBS of the left lung were performed, red blood cells lysed and total inflammatory cells counted, cytospun, air dried and stained with May-Grunwald-Giemsa for differential counts. For histological analysis, lung tissue was perfused with saline *via* cardiac puncture, inflated (500µL) and fixed in formalin prior to mounting, sectioning and staining. Hematoxylin and eosin-stained lung sections were used to assess parenchymal inflammation by counting the number of inflammatory cells in 10 randomised fields of view at 100x magnification,^18, 20, 21^ and emphysema-like alveolar enlargement using the mean linear intercept, with assessor blinded to sample identity.^20, 22^

**Lung function**

Mice were anaesthetized with ketamine (100 mg/kg) and xylazine (10 mg/kg). Tracheotomy was performed, and mice cannulated. A BioSystem Forced Maneuvers (Buxco, Wilminton, NC) and a flexivent apparatus (Legacy System, SCIREQ, Montreal, Canada) were used to measure all lung function parameters. Each perturbation was conducted a minimum of three times and the average calculated for each mouse.^19-24^

**Colon histopathology**

For histological analysis, colon and ileum tissue was fixed in formalin prior to mounting, sectioning, and staining with H&E. Images were taken at 20x magnification and sub-mucosal fragmentation area measured using ImageJ software. Blood vessels were enumerated along the entire colon and corrected for sub-mucosal length.^22^

**RNA extraction, cDNA synthesis, and qPCR analysis**

Colon tissues were snap frozen and stored at 80°C. RNA was extracted using a previously described TRIzol (ThermoFisher Scientific) protocol utilizing phase separation in chloroform and RNA precipitation in isopropanol.^19-21, 25^ Random-primed reverse transcriptions were performed followed by real-time qPCRs with SYBR-green–based detection using a Viia 7 Real Time PCR system.^19-21, 25^ Specific mRNA transcripts were assessed and expressed as relative abundance compared to β-actin (*Actb)* expression (Table S16).

**CyTOF analysis**

For time-of-flight mass cytometry, whole blood was collected into ethylenediaminetetraacetic acid (EDTA)-coated tubes and spleen tissue disaggregated using a 70µm cell strainer. Bone marrow was collected and immediately incubated with the cell cycle marker iododeoxyuridine (IdU; 10µM, 1hr, 37°C). For all samples, red blood cells were lysed and cells quantified using trypan blue. Single-cell suspensions were stained with the viability marker cisplatin (5µM, 5 minutes, room temperature), quenched (5% foetal calf serum, 5mM EDTA in PBS) and collected by centrifugation (300x*g*, 3 minutes, 4°C). Fc receptors were blocked with metal conjugated anti-mouse CD16/32 antibody (Table S17, 30 min, 4°C) and quenched before staining with surface antibodies (Table S17, 30 min, 4°C). Cells were washed and fixed (4% paraformaldehyde, 4°C overnight) before permeabilization in methanol (10 minutes, 4°C), washing and staining with intracellular antibodies (Table S17, 45 minutes, room temperature). Samples were washed thrice in permeabilization buffer (Thermo Fisher) and stored in 4% paraformaldehyde at 4°C until analysis using a Helios instrument (Fluidigm).

The expression levels of 38 markers (Table S17) measured by CyTOF was used as input for downstream analysis. Before analysis, a maximum of 10,000 cells for each sample were randomly selected. Protein expression was transformed with a logicle transformation^26^ (width=0.25, top=16,409, full width=4.5) and used for Uniform Manifold Approximation and Projection for Dimension Reduction (UMAP) calculated using the naïve R implementation (umap 0.2.7) of the algorithm with the following settings: number of neighbors=15, number of components=2, distance metric=Euclidean, minimum distance=0.2. Cells clusters showing similar marker expression were calculated according to the Phenograph algorithm (Rphenograph, Github JinmiaoChenLab v. 0.99.1)^27^ with standard setting and number of nearest neighbors (k)=60. Data were visualized according to the coordinate on the UMAP dimensions and the cluster assignment of each cell. Confusion matrices were calculated normalizing the number of cells in each experimental group to 1,000 and calculating the contribution of each condition in each cluster to avoid bias from different cell numbers in experimental groups. Finally, a heatmap was generated using the mean marker expression in each cluster scaled for visualization. Data were visualized using the R packages ggplot2 (v. 3.3.2) and pheatmap (v. 1.0.12). All analysis was performed in a dockerized environment based on R 4.0.3 and Bioconductor 3.12 (lorenzobonaguro/flowtools:v2).

**Cell culture**

Raw 264.7 monocytes (1x10^6^) were seeded into 12-well plates and allowed to adhere overnight. Feces from CS- or air-exposed mice were homogenized in DMEM (100mg/mL) and sterile-filtered. Cells were incubated in media alone, or 1:1,000 dilutions of filtered fecal homogenate for 24 hours. During the final 4 hours of incubation lipopolysaccharide (1µg/mL), monophosphoryl lipid A (4µg/mL), lipoteichoic acid (1µg/mL) or dimethyl sulfoxide (0.5%) were added to cell media. Media was collected and stored at -80°C until TNFα protein was assessed using Duoset ELISA kits (R & D Systems).

**DNA extraction and 16S rRNA amplicon community profiling**

DNA was extracted from fecal samples using the PowerSoil DNA extraction kit according to the manufacturer’s instructions (MoBio, Carlsbad, California). For community profiling, the V5-V8 region of the 16S rRNA gene was amplified using a modified two-step PCR protocol.^28^ The first round 50μl reaction mixture comprised 20ng of template DNA, molecular biology grade water, 5μl 10x PCR buffer (Fisher Scientific), 4μl of 25mM MgCl2 (Fisher Scientific), 1.5μl BSA (New England Biolabs), 1μl of 10mM dNTPmix (Fisher Scientific), 1μl of 10nM forward primer 803F (803Fa-5’: TTAGATACCCTGGTAGTC; 803Fb-5’: TTAGATACCCSGGTAGTC; 803Fc-5’: TTAGATACCCYHGTAGTC; 803Fd-5’: TTAGAGACCCYGGTAGTC; mixed at ratios of (2a:b:c:d), 1 μl of 10nM reverse primer 1392wR (ACGGGCGGTGWGTRC)) and 0.2μl of 5U/ul Taq Polymerase (Fisher Scientific). Cycling conditions were as follows: 95°C for 3 minutes, 30 cycles of 95°C for 30 seconds, 55°C for 30 seconds and 74°C for 30 seconds followed by a final extension of 74°C for 10 minutes. PCR products were re-amplified in a second reaction using primers containing (5’-3’) 454 adaptor sequences, barcode (only for reverse primer), linker sequence and the gene-specific primer sequence (803F and 1392wR: pyroLSSU803F – CCTATCCCCTGTGTGCCTTGGCAGTCTCAGTTAGAKACCCBNGTAGTC

pyroLSSU1392wR - CCATCTCATCCCTGCGTGTCTCCGACTCAGACATAGTACGGGCGGTGWGTRC). For the second step reactions, 2 μl of the first step PCR product was added as template to a 50 μl reaction. PCR conditions were the same as those for the first step with the number of cycles reduced to 10. Amplicons were purified using AMPure beads (Beckman Coulter) and pooled in equal ratios. Amplicon pools were sequenced from the reverse primer using 454 GS-FLX Titanium chemistry.

Amplicon sequences were demultiplexed and converted to fastq files using QIIME v1.9.1 scripts split_libraries.py, convert_fastaqual_fastq.py and split_sequence_file_on_sample_ids.py. Reads were then quality filtered and trimmed of adapter sequence using BBMap v38.41 (<https://sourceforge.net/projects/bbmap/> via BBDuk; ktrim=l k=15 mink=10 hdist=1 interleaved=f copyundefined=t mm=f qtrim=rl trimq=10 minlen=400). Sequence variants were defined using QIIME2 v2019.10^29^ and DADA2 (via denoise-pyro).^30^ Taxonomy was assigned using QIIME2 feature-classifier ^31^ classify-consensus-blast against the Silva v138 database.^32^

**Proteomics of mouse feces**

Feces (100µg) were added to 500µL of sodium bicarbonate, vortexed with 2.5-mm glass beads and supernatants collected (300x*g*, 5 min, 4°C). Pellets were resuspended in sodium bicarbonate, centrifuged and supernatants collected a further two times. Supernatants were combined and washes repeated to remove debris before final cell collection (14,000x*g*, 20 min, 4°C). Cells were aliquoted (~30mg wet weight per aliquot) and subjected to enzymatic degradation (20µL lysozyme in buffer containing 20mM TrisHCl, 2 mM EDTA and 1% Triton X-100; 30 min, 37°C). Incubation was halted with sodium bicarbonate and cell debris removed by centrifugation (12,000x*g*, 20 min, 4°C). Proteins (150µg) were precipitated, resuspended in 8M urea, reduced with dithiothreitol and alkylated with iodoacetamide. Samples were diluted in 50mM ammonium bicarbonate to a final urea concentration of 1M and digested overnight with trypsin (50:1 protein:enzyme, 37°C). Following acidification in 1% trifluoroacetic acid, 20μg of peptides were dried by vacuum concentration, resuspended in 2% acetonitrile/0.1% trifluoroacetic acid and 500ng injected onto a trapping column for pre-concentration (Acclaim Pepmap100, 20x0.075mm, 3μm C18, Thermo Scientific), followed by nanoflow liquid chromatography (Ultimate 3000RSLCnano, Thermo Scientific). Peptide separation was achieved over a 250x0.075 mm ID, PepMap 2µm EasySpray C18 column using the following mobile phases: water, 0.1% formic acid (solvent A) and 80% acetonitrile incorporating 0.1% formic acid (solvent B). Peptides were resolved using a linear gradient from 2% B to 35% B over 70 min at a constant flow of 300nl/min. Peptides were introduced via an EasySpray Nano source coupled to a Q-Exactive Plus Quadrupole Orbitrap mass spectrometer (Thermo Scientific) and subjected to data dependent tandem mass spectrometry (MS/MS) using the following parameters. Full MS scans were acquired in profile mode, positive polarity at a resolution of 70,000, automatic gain control target 1x10^6^ and a max injection time of 50ms. Data dependent MS/MS was then preformed at a resolution of 17,500 on the top 20 ions with a charge ≥ 2, automatic gain control target of 5x10^5^, max injection time of 120ms, using a stepped NCE of 26 and 30. Dynamic exclusion was set at 15 seconds. .raw files were processed in Proteome Discoverer v2.1 using the Sequest HT algorithm ^33^ and searched against the mouse gut microbiota GigaDB database.^34^

**Integration of proteomics and metagenomics**

Metagenomics results were collapsed by adding relative abundances of bacteria from the same genera. Genera and proteins present in ≥50% of samples were included for analysis. Both microbiome and proteomic data were integrated through correlation analysis using the psych package in R^35^ and Diablo mixOmics package.

**Metabolomics**

Caecum contents were collected from mice exposed to CS or normal air for 12 weeks, snap frozen and shipped on dry ice to Metabolon Inc (Durham, NC). Samples were analysed using the Global HD4 mass spectrometry platform using ultra-high performance liquid chromatography. Peaks were identified by comparison to a library of purified standards (or recurrent unknown compounds) containing retention time/index, mass:charge ratio (m/z), and chromatographic data (including MS/MS spectral data), and the relative abundance of each metabolite scaled such that the median value was equal to 1.
